# Optimizing a High Performing Multiplex-CRISPRi *P. putida* Strain with Integrated Metabolomics and ^13^C-Metabolic Flux Analyses

**DOI:** 10.1101/2021.12.23.473729

**Authors:** Jeffrey J. Czajka, Deepanwita Banerjee, Thomas Eng, Javier Menasalvas, Chunsheng Yan, Nathalie Munoz Munoz, Brenton C. Poirier, Young-Mo Kim, Scott E. Baker, Yinjie J. Tang, Aindrila Mukhopadhyay

**Affiliations:** Department of Energy, Environmental and Chemical Engineering, Washington University, St. Louis, MO, 63130, USA; Joint BioEnergy Institute, Lawrence Berkeley National Laboratory, Emeryville, CA, 94608, USA; Biological Systems and Engineering Division, Lawrence Berkeley National Laboratory, Berkeley, CA, 94720, USA; Earth and Biological Sciences Directorate, Pacific Northwest National Laboratory, Richland, WA, 99352, USA; Agile BioFoundry, Emeryville, CA 94608, USA; QB3 Institute, University of California, Berkeley, CA 94720, USA; Environmental Genomics and Systems Biology Division, Lawrence Berkeley National Laboratory, Berkeley, CA, 94720, USA

**Keywords:** PHA, ^13^C Metabolic flux analysis, *Pseudomonas putida* KT2440, Multiplexed CRISPR interference, Recombineering, Indigoidine

## Abstract

Microbial cell factory development often faces bottlenecks after initial rounds of design-build-test-learn (DBTL) cycles as engineered producers respond unpredictably to further genetic modifications. Thus, deciphering metabolic flux and correcting bottlenecks are key components of DBTL cycles. Here, a 14-gene edited *Pseudomonas putida* KT2440 strain for heterologous indigoidine production was examined using both ^13^C–metabolic flux analysis (^13^C–MFA) and metabolite measurements. The results indicated the conservation of the cyclic Entner-Doudoroff (ED)-EMP pathway flux, downregulation of the TCA cycle and pyruvate shunt, and glyoxylate shunt activation. At the metabolite level, the CRISPR/dCpf1-interference mediated multiplex repression decreased gluconate/2-ketogluconate secretion and altered several intracellular TCA metabolite concentrations, leading to succinate overflow. Further strain engineering based on the metabolic knowledge first employed an optimal ribosome binding site (RBS) to achieve stronger product-substrate growth coupling (1.6–fold increase). Then, deletion strains were constructed using ssDNA recombineering. Of the five strains tested, deletion of the PHA operon (Δ*phaAZC-IID*) resulted in a 2.2–fold increase in growth phase production compared to the optimal RBS construct. After 72 h of batch cultivation, the Δ*phaAZC-IID* strain had 1.5–fold and 1.8–fold increases of indigoidine titer compared to the improved RBS construct and the original strain, respectively. Overall, the findings provided actionable DBTL targets as well as insights into physiological responses and flux buffering when new recombineering tools were used for engineering *P. putida* KT2440.

## 1. Introduction

*Pseudomonas putida* KT2440 is emerging as an advantageous metabolic engineering chassis due to its genetic tractability, rapid growth rate, and robust ability to grow on lignin and non-lignin derived feedstocks (Nikel and de Lorenzo, 2018). Previous efforts have targeted natural and heterologous bioproducts for expression in *P. putida* including biofuels (phenazine, methyl ketones)(Askitosari et al., 2019; Dong et al., 2019), lipids (rhamnolipid)(Arnold et al., 2019), polymers (polyhydroxyalkanoate)(Yang et al., 2019), and organic acids (adipic acid) (Niu et al., 2020). Recently, the production of the non-ribosomal peptide indigoidine in *P. putida* KT2440 achieved promising titers of ~2 g/L in shaking flasks cultivations and ~26 g/L during fed-batch cultivations (Banerjee et al., 2020). This previous study genomically integrated the two heterologous genes (*bpsA* and *sfp*) required to catalyze the conversion of glutamine to indigoidine and implemented a complex single design-build-test portion of a design-build-test-learn (DBTL) cycle to improve production. The design was generated using the constrained minimal cut set (cMCS) genome-scale modeling technique which identifies genetic targets for deletion to obtain strong product-substrate growth coupling (Klamt and Mahadevan, 2015; Trinh et al., 2009). A multiplexed CRISPR-interference (CRISPRi) system was then used to simultaneously knock-down 14 predicted gene targets. The product-substrate paired growth-coupling strain (PSP strain) had a 30% improvement in indigoidine production compared to the base strain despite the partial design implementation (Banerjee et al., 2020). This PSP strain provided a valuable system to examine a highly engineered strain and to investigate potential emergent metabolic features and phenotypes. Furthermore, advanced DBTL cycles can struggle to achieve the same level of improvements in initial cycles due to more failure routes in strains. Insight gained here on the engineered PSP strain’s metabolism can guide further strain designs and help increase the effectiveness of a second DBTL cycle.

^13^C-Metabolic flux analysis (MFA) is a technique that measures intracellular enzymatic rates and has been employed to decipher cellular energy metabolism (He et al., 2014; You et al., 2015), responses to genetic perturbation (Long et al., 2016), and pathway regulation (Long and Antoniewicz, 2019). Previous ^13^C–MFA studies of *P. putida* metabolism during growth on glucose revealed several core metabolic features, such as the Entner-Doudoroff (ED)-EMP cycle, an active pyruvate shunt, and an inactive glyoxylate shunt (Kohlstedt and Wittmann, 2019; Nikel et al., 2021, 2015). However, the metabolic rewiring of *P. putida* strains in response to complex editing of central pathways remains poorly understood. The indigoidine producing *P. putida* strains were an ideal system to investigate as production was previously characterized in a minimal defined medium compatible with ^13^C-MFA (Banerjee et al., 2020). Thus, this study aimed to characterize the metabolic responses with direct measurements of cellular output via fluxomic data. Metabolomic data was also collected as it is essential to combine MFA with targeted metabolite analyses to precisely identify congestion nodes (bottlenecks) in the flux network (Cocuron et al., 2019; Czajka et al., 2020a; Raamsdonk et al., 2001). This study integrated multi-level omics information to decipher metabolic rewiring and to select promising gene deletion targets for improved production - which in turn were implemented via high performance recombineering techniques.

## 2. Materials and Methods

### 2.1 Strains and plasmids

All strains used in this study are listed in Table 1. The strains analyzed via ^13^C–MFA were the wildtype (WT), the strain containing engineered indigoidine production pathway (Eng), and the strain containing both the engineered indigoidine production pathway and the CRISPRi product-substrate pairing plasmid (PSP).

**Table 1.**
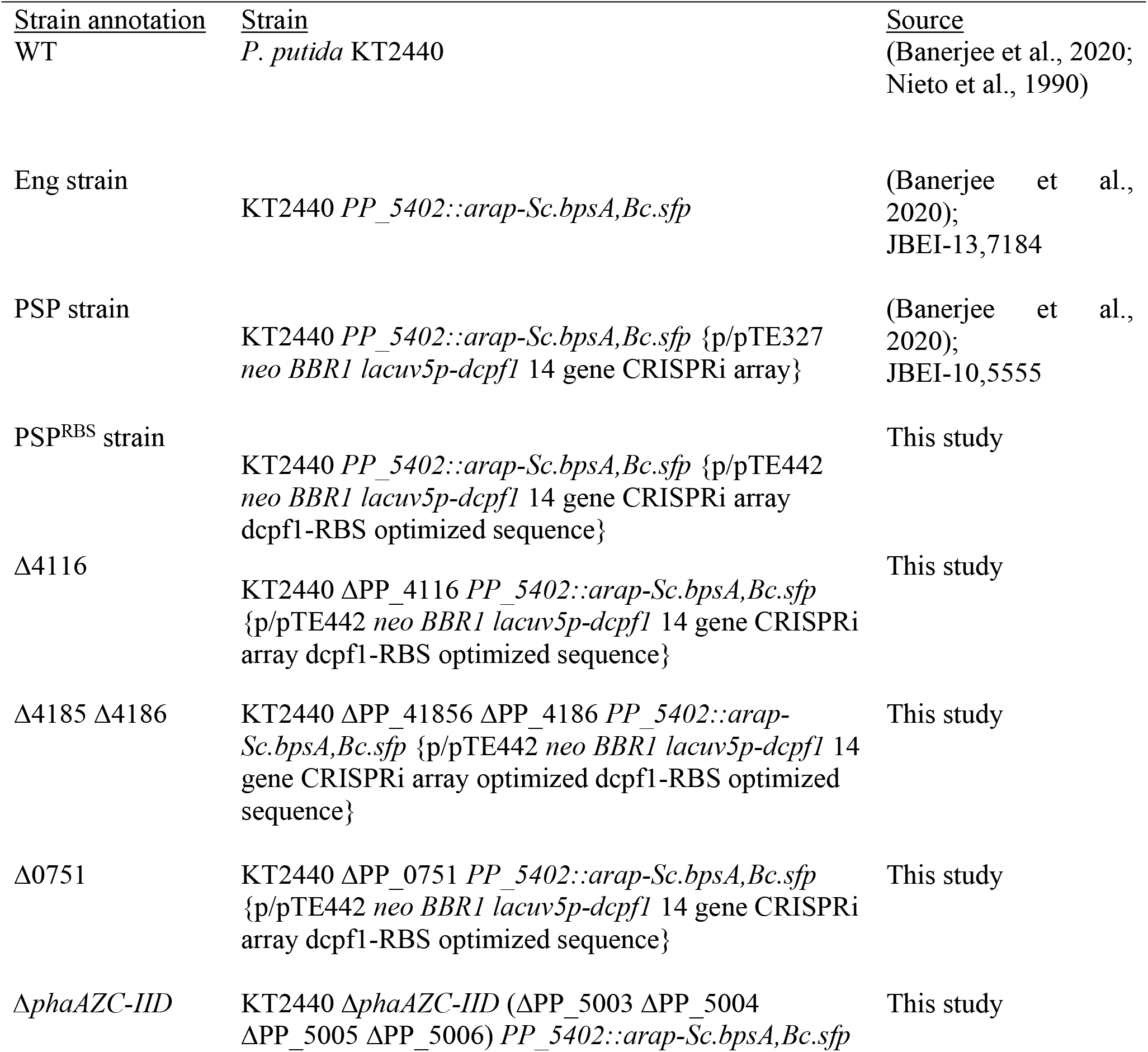
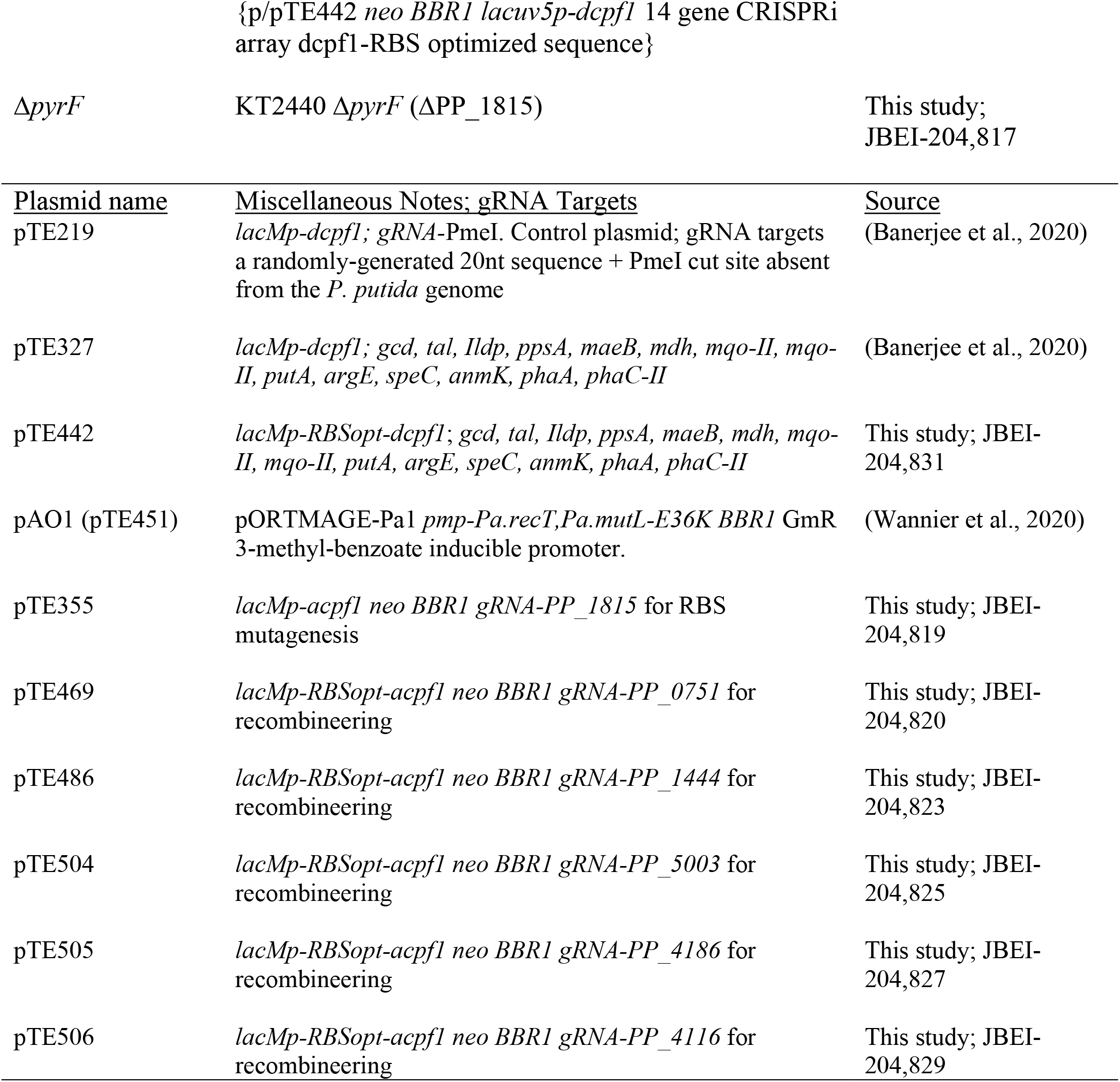
Strains and plasmids used in this study.

Expression of the heterologous indigoidine pathway is under the control of an arabinose inducible promoter (Banerjee et al., 2020). The PSP strain gRNA expression cassette is induced by IPTG. The gRNA targeted 14 genes for downregulation (*gcd, tal, Ildp, ppsA, maeB, mdh, mqo-II, mqo-II, putA, argE, speC, anmK, phaA, phaC-II*). All plasmids contain *neo* which confers resistance to kanamycin for selection. The endonuclease inactive allele of *Francisella novicida U112 cpf1* (cpf1-D917N) is described as dcpf1. The endonuclease active allele is referred to as acpf1. The sequences of the plasmids generated in this study may be visualized at public-registry.jbei.org.

### 2.2 Chemicals and Growth Medium

Labeled substrates [1,2-^13^C glucose, 6-^13^C glucose, 1-^13^C glucose, U-^13^C_6_ glucose] were purchased from Omicron Biochemicals (South Bend, IN) or Sigma-Aldrich (St. Louis, MO). All other chemicals were purchased from Sigma-Aldrich. *P. putida* strains were grown in LB medium or M9 minimal medium [per liter, 2 g (NH_4_)_2_SO_4_, 6.8 g Na_2_HPO_4_, 3 g KH_2_PO_4_, 0.5 g NaCl, 1 mL trace elements solution (Teknova, Hollister, CA), 100 μL 1 M CaCl_2_, 2 mL 1 M MgSO_4_] supplemented with 10 g/L of glucose. Arabinose (3 g/L), IPTG (0.5 mM), and Kanamycin (50 μg/L) were added as necessary for indigoidine production.

### 2.3 Cell Cultivation

*P. putida* cultivated for metabolomic and fluxomic analyses were grown in 14 mL of liquid volume in 50 mL unbaffled shaking flasks at 30 °C and 200 rpm. Seed cultures were inoculated from fresh plates (<3 days old) in 5 mL of LB medium and grown overnight. A 1.4% inoculation ratio was used to start 14 mL liquid cultures of M9 minimal medium. Cells were subcultured in M9 production medium at an initial OD_600_ of 0.06 – 0.10 and samples for the metabolomic, fluxomic, and production assay experiments were collected from the subculture. Fluxomic experiments substituted the unlabeled glucose in the subculture with either 1,2-^13^C glucose, a 80:20 mixture of 6-^13^C glucose:U-^13^C_6_ glucose or a 80:20 mixture of 1-^13^C glucose:U-^13^C_6_ glucose (Cambridge Isotope, MA and Omicron Biochemicals, IN). Cultures used as internal standards for metabolomic measurements were grown in two subsequent M9 minimal mediums cultures containing 100% U-^13^C_6_.

### 2.4 Dry Cell Weight Measurements

The dry cell weight per OD_600_ measurements were collected from cultures grown with M9 minimal medium (1^st^ subculture). Cells were inoculated to an approximate OD_600_ of 0.06 in 50 mL of M9 minimal medium in 250 unbaffled shaking flasks. OD_600_ was measured throughout the growth with biological replicates harvested at various OD_600_ values via centrifugation at 5,000 x g for 10 mins. The supernatant was then discarded, the pellet washed with 0.9% NaCl, and the liquid re-centrifuged. Samples were frozen at −80 °C before lyophilization and measurement. One OD_600_ unit was 0.37 ± 0.02 g/L biomass during the growth phase (**Fig. S1**).

### 2.5 Sample Collection and Processing for Metabolomic Analysis

Metabolomic samples were collected at both the growth phase (OD_600_ range of 0.8 – 1) and the production phase (24 hrs, OD_600_ range of 8 – 10). The sampling process involved rapidly quenching metabolism using a carbon-free medium in a liquid nitrogen bath. Specifically, the culture was poured into a chilled (~0 °C) M9 medium solution that was rapidly stirred in a liquid nitrogen bath until the culture reached ~0 °C. Cells were then pelleted at 5,000 x g for 5 mins (at 1 °C), flash frozen with liquid nitrogen, and stored at −80 °C until metabolite extractions were performed. Intracellular metabolites were extracted in a 1 mL 7:3 MeOH:Chloroform solution at 4 °C, and then processed and analyzed as previously described (Czajka et al., 2020a). Intracellular concentrations were normalized via the dry cell weight correlations (**Fig. S1**). Cultivation medium was filtered through 0.2 μm sterile filters and lyophilized for extracellular metabolite measurements using the same method. A second set of extracellular metabolites were quantified from 20 μl of spent medium that were dried under vacuum. Chemical derivatization, analysis by GC-MS (same instrument as citation), and data processing was done as previously described (Pomraning et al., 2021). Glucose content was determined via enzymatic kit (R-Biopham, Darmstadt, Germany) per manufacturer’s instructions.

### 2.6 Sample Collection and Processing for Proteinogenic Measurements

Proteinogenic amino acid label incorporation samples were harvested from cultures that were grown to an OD_600_ range of 0.7 – 1.1. Cells were pelleted at 5,000 x g for 5 mins, the supernatant was discarded, and the pellet was frozen at −80 °C until processing. Proteins from cultures grown with 1-^13^C or 6-1^3^C glucose tracers were collected from a MPLEx extraction (Nakayasu et al., 2016), hydrolyzed with 6 N HCl at 100°C for 20 hours and dried with a speed vacuum concentrator. The amino acids were then dissolved in 20 μL of pure pyridine and chemically derivatized using 80 μL of tert-butyldimethylsilyl trifluoromethanesulfonate (TBDMS) at 70 °C for one hour. The raw data were analyzed and amino acid fragments were corrected for natural labeling abundance by the software DExSI (Dagley and McConville, 2018). All other samples were hydrolyzed with 1 mL of 6 N HCl at 100 °C for 20 h, dried with filtered air, and derivatized using 100 μL of TBDMS in 100 μL of THF at 70 °C for one hour. All amino acid derivatized samples were analyzed via GC-MS equipped with a HP-5MS column as previously described (Hollinshead et al., 2019). The amino acid fragments were corrected for natural labeling abundance according to the published method (Wahl et al., 2004).

### 2.7 Indigoidine measurements

Indigoidine production and quantification was performed as previously described with slight modifications (Banerjee et al., 2020). Briefly, either 500 μL (OD_600_ = 1) or 100 μL (24 h, 48 h samples) of liquid culture was pelleted at 24,000 x g for 2 min. The supernatant was discarded, and 500 μL of dimethylsulfoxide (DMSO) was used to resuspend and extract the indigoidine via vortexing (10 mins). Additional DMSO was added if the pellet was not fully dissolved. Absorption was measured at 612 nm in a Cary 60 UV-Vis Spectrometer (Agilent Technologies). Levels of indigoidine are reported as a function of absorbance with the previously determined calibration curve used to estimate titers (Banerjee et al., 2020).

### 2.8 Flux Modeling

A core *P. putida* metabolic network was constructed from published resources (Kohlstedt and Wittmann, 2019). The INCA software package was used to analyze the metabolic network (Young, 2014) for parallel tracer experiments. The WUflux software was utilized to cross-validate the flux calculations using the 1,2-^13^C glucose derived data (He et al., 2016). The average growth rates, indigoidine production rates, and secretion rates were used to constrain the network to generate customized MFA models that represented WT, Eng, and the PSP strains. The glucose uptake measurements and biomass formation at 6 h were used to calculate the uptake per biomass yield and to further constrain the models (**Fig. S2**). The *P. putida* biomass equation utilized here was previously reported and derived from biomass yield and represents the normalized precursor drainage to obtain the experimental growth rates (Kohlstedt and Wittmann, 2019). The phosphoenolpyruvate carboxylase reaction has been annotated as a pseudogene in *P. putida* (Nelson et al., 2002; Nikel et al., 2021). Inclusion of the reaction did not result in changes to the fit, thus the reaction was included to function in the direction annotated within the KEGG database (https://www.kegg.jp/pathway/ppu00620+K01595).

### 2.9 Identification of an Optimized RBS Sequence for Cpf1 Function

A small library of ribosome binding site variants for *Francisella novicida U112 cpf1* were calculated using denovoDNA (Salis et al., 2009) and incorporated into a plasmid containing an endonuclease active (D917) allele of *cpf1* and a gRNA targeting *pyrF* (PP_1815) using site directed mutagenesis (Deng and Nickoloff, 1992) with Q5 polymerase (NEB). The RBS mutant library was encoded by the degenerate sequence 5’-GYAGAASAKTCMAAATGGKGASRTGGAT-3’. The RBS library was transformed into *E. coli* DH10-beta competent cells (NEB); approximately 50 single colonies were picked, inoculated into liquid LB media with 50 μg/mL Kanamycin, and single plasmids were extracted using a Miniprep plasmid DNA extraction kit (Qiagen Research, Germantown, MA). *P. putida* KT2440 and the Δ*pyrF* strain were made electrocompetent (Wang et al., 2009) and transformed with a variant plasmid from the RBS library. Following an hour outgrowth in LB media at 30 °C (200 rpm shaking), the entirety of the transformation was spun down and plated on an LB Kan plate and incubated at 30 °C overnight. Candidate RBS-variant *cpf1* plasmids were identified by the following criteria: >200 CFUs/μg plasmid DNA in the Δ*pyrF* strain background; <5 CFUs/μg plasmid DNA in the KT2440 wild-type background. One candidate clone, RBS isolate number 30, met this criterion. The RBS sequence was identified by Sanger sequencing and corresponded to the following sequence identity: 5’-GCAGAACAGTCAAAATGGGGACGTGGAT-3’. This RBS sequence was introduced into the multiplex CRISPRi/dCpf1 plasmid pTE327 using site directed mutagenesis between the J2113 promoter and start codon of *cpf1* and subsequently was given the accession ID pTE442.

### 2.10 Generation of Deletion Strains via Recombineering & Cfp1/CRISPR Selection

Deletion mutants in this study were generated using a RecT-family recombinase following a modified protocol based on (Aparicio et al., 2020). Briefly, fresh transformants harboring pTE451/pAO1 were selected using LB agar plates supplemented with 30 μg/mL gentamicin. Single colonies from fresh plates (<5 days old) were used to inoculate LB gentamicin liquid cultures and grown overnight at 30 °C (200 rpm shaking). 2.5 mL of the overnight culture (OD_600_= ~4) was used to inoculate 25 mLs of fresh LB gentamicin medium in a 250 mL baffled shaking flask and grown for 1 h at 30 °C and 200 rpm. Afterwards, recombinase expression was induced with 3-methyl-benzoate (Sigma: T36609; M-Toluic Acid 99% purity) at final concentration of 1 mM. Cells were incubated for 30 min after induction with shaking, decanted into a 50 mL falcon tube, and centrifuged for 5 mins at 3,000 x *g* at 4 °C. The cell pellet was resuspended with 10% glycerol and transferred into an Eppendorf tube, washed with glycerol three additional times, and resuspended into a final volume of 1 mL of 10% glycerol.

For each recombineering event, 50 μL of the aliquoted cells were mixed with 1 μL of the single stranded oligonucleotide (2 μM final concentration) (**Table S1**) and 50 ng of the appropriate cpf1-gRNA (endonuclease active) plasmid (**Table 1**). The above-mentioned components were electroporated into *P. putida* using 2 mm-gap cuvettes and the Bio-Rad MicroPulser (program EC2 - 2.5 kV). After pulsing, cells were immediately recovered in 600 μL of Terrific Broth (TB) and incubated for 3 h at 30 °C, 1,000 rpm in a bench-top thermomixer. After the outgrowth phase, 250 μL of the suspension was plated on solid agar LB kanamycin plates for 1-3 days at 30 °C. Colonies were pinpoint-sized after 24 h but were readily visualized after 48 h.

Between 8–30 clones were selected for genotyping by colony PCR using NEB OneTaq Quick-Load 2X Master Mix with Standard Buffer (catalog # M0486L) following the manufacturer’s protocol. Before PCR, colony biomass was boiled at 94 °C for 45 minutes in 50 μL 20 mM NaOH. 2 μL of the boiled solution was used in a 25 μL PCR reaction. The annealing temperature was calculated using the webtool, NEB Tm calculator (tmcalculator.neb.com, New England Biolabs, Ipswitch, MA). The loci targeted for deletion were genotyped using colony PCR with primers that bind to the 5’ upstream and 3’ downstream regions adjacent to the targeted open reading frame (**Table S5**). PCR products were analyzed using standard techniques for agarose gel electrophoresis. After genotyping, both the recombineering plasmid and CRISPR plasmid were cured from the mutant by allowing random segregation; sensitivity to either antibiotic was verified by patching clones to media containing either antibiotic.

## 3. Results

### 3.1 Strain Physiology and Production Characteristics

The first step in performing ^13^C–MFA was to evaluate the cell physiological state under the desired cultivation conditions. It was previously reported that the strain containing the production pathway and the CRISPRi growth-coupling plasmid (PSP strain, **Table 1**) maintained a desirable phenotype under a variety of scales (deep-well microplates, shaking flasks, and bioreactors) (Banerjee et al., 2020). To minimize labeled substrate use, the culture volume was scaled down to a 14 mL fill volume in 50 mL unbaffled shaking flasks (**Fig. S2**). The PSP strain showed a ~11% increase in production compared to the Eng strain across several runs in the 14 mL format (**Fig. 1**), consistent with previously reported titers (Banerjee et al., 2020). The maximum growth rates of the three strains in 14 mL cultures were not statistically different, although there was a longer lag phase for the PSP strain due to the addition of antibiotics necessary to retain the CRISPR interference (CRISPRi) plasmid (**Fig. 1**). This increased lag did not cause a decrease in the biomass yield coefficient of the PSP strain, with yield coefficients in the physiological range of previous reports (del Castillo et al., 2007; Kohlstedt and Wittmann, 2019) (0.45 ± 0.02 g biomass/g glucose and 0.44 ± 0.04 g biomass/g glucose consumed for the Eng and PSP strains, respectively).

**Figure 1.**
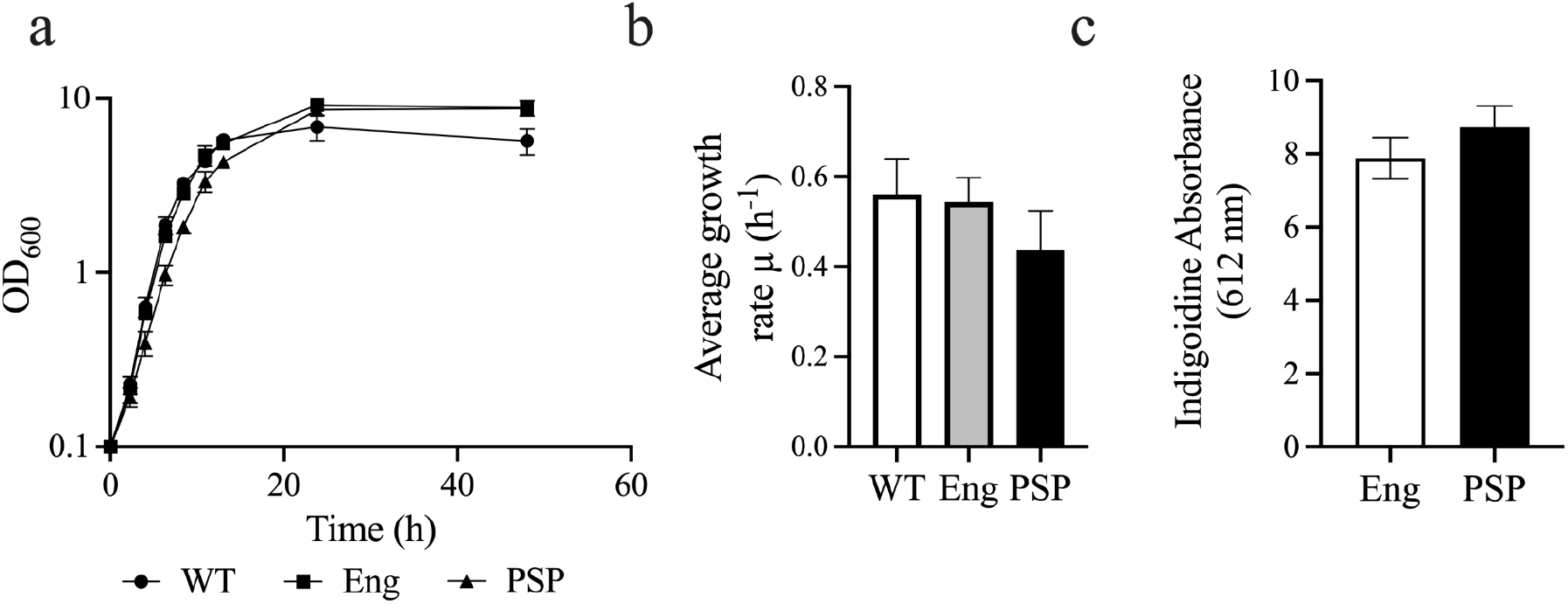
Growth characterization of wild type *P. putida* and indigoidine producing strains. (a) Growth curves (log scale) on glucose in M9 minimal medium (n=4). The PSP strain was significantly (p-value <0.05) different from the WT and Eng strains. (b) Average growth rates (n=14). Error bars represent the standard error. (c) Indigoidine production at 48 h (n=6). The indigoidine absorbance for the PSP strain is ~1.8 ± 0.3 g/L using a previously reported standard curve (Banerjee et al., 2020). Error bars represent the standard error.

The next step to employ ^13^C–MFA was to determine the extracellular metabolites secretion rates to constrain the MFA model. *P. putida* has been previously reported to secrete gluconate and 2-ketogluconate (2KG) during growth on glucose minimal medium (del Castillo et al., 2007; Kohlstedt and Wittmann, 2019; Nikel et al., 2021, 2015) or complex LB medium (Molina et al., 2019a, 2019b). An analysis of the extracellular medium revealed both compounds were secreted by the WT strain, reaching of 1.4 ± 0.1 mM of gluconate and 3.9 ± 0.3 mM of 2KG (~1.0 ± 0.1 g/L total secretion) towards the end of the growth phase (**Fig. 2a**). Interestingly, 2KG was secreted in higher quantities than gluconate under the cultivation conditions tested, in contrast to the previous minimal medium studies but in agreement with the LB studies, indicating periplasm oxidation may be sensitive to cultivation conditions. The Eng strain had increased secretion of 2KG (5.4 ± 0.9 mM) and gluconate (1.8 ± 0.3 mM) compared to the WT, while expression of the CRISPRi construct successfully reduced the amount of secreted compounds and shifted the excretion towards an equal amount of both compounds (1.4 ± 0.1 mM 2KG and 1.6 ± 0.1 mM gluconate). Both compounds were re-consumed by the 24 h mark, with the exception of a small amount of remaining gluconate in the PSP strain (**Fig. 2b**). Several minor byproducts were present in all strains; however, going into the production phase, the PSP strain secreted several organic acids not observed in the other strains including pyruvate, lactate, and succinate (**Fig. S3**).

**Figure 2.**
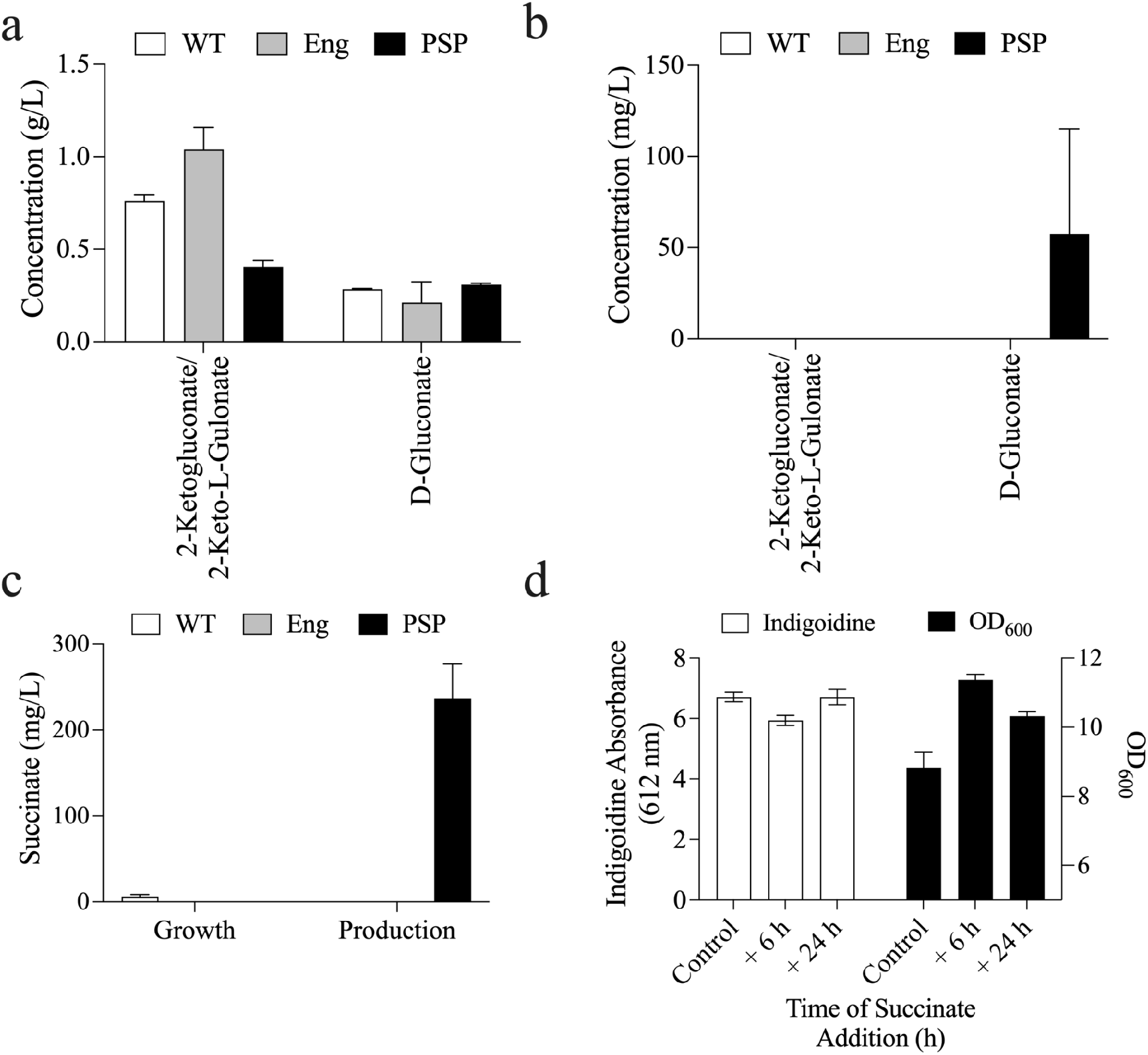
Succinate dynamics and indigoidine production. (a) Secreted extracellular sugars during the growth phase. (b) Extracellular sugars during the production phase. (c) Succinate overflow during the growth (6 h) and production (24 h) phases. (d) Indigoidine production after succinate addition during either 6 or 24 h after inoculation. The indigoidine absorbance for the control is ~1.4 ± 0.1 g/L using a previously reported standard curve (Banerjee et al., 2020). Error bars represent the standard error (a,b n=2; c, n=3; d, n = 4 (control), n = 3 (6 h and 24 h time points).

Unexpectedly, further analysis revealed that succinate accumulated as a relatively large byproduct with 240 ± 40 mg/L detected in the production phase (**Fig. 2c**). The previous cMCS-based modeling (Banerjee et al., 2020) used for designing the PSP strain had not assumed an overflow of succinate, as *P. putida* succinate secretion was previously reported only under nitrogen-limiting conditions during growth on glycerol (Beckers et al., 2016). Additionally, the modeling predicted succinate to be an incompatible substrate for production (i.e., succinate would not be sufficient as a carbon source to generate precursors for both biomass growth and indigoidine production (Banerjee et al., 2020)(**Fig. S4**)). To verify the cMCS prediction, succinate was added to cultivations of the Eng strain (3 g/L) at either 6 h or 24 h. The additional carbon led to increased biomass accumulation but not increased indigoidine levels (**Fig. 2d**). Therefore, the secreted succinate represented a loss of carbon from the system that was not being directed towards product synthesis. After collecting the necessary information to constrain the core *P. putida* model, ^13^C-MFA was performed.

### 3.2 Steady-State Flux Modeling

Parallel ^13^C–MFA tracer experiments were performed using three different labeled glucose mixtures as the carbon source (each tracer was used in an independent culture, **see Methods Section 2.3**). Based on previously established methods, at least three doublings of growth were allowed to occur in the labeled medium after inoculation before biomass was harvested (Wiechert et al., 2007). Steady-state flux fitting requires an assumption of metabolic and isotopic steady-state. The assumption was verified by growing the cells on a mixture of labeled glucose substrates and harvesting the biomass at different points within the desired range of growth. The resulting amino acid labeling experiment profiles were similar, with only small, non-significant differences (p-values <0.05, student’s t-test) observed between timepoints (**Fig. S5**). After verification of the steady-state assumption, ^13^C-MFA labeling experiments were performed.

The fluxes were simulated with the appropriate carbon transitions of central carbon metabolism constructed based on previously published work (Kohlstedt & Wittmann, 2019), **see Methods Section 2.8**). As small metabolite labeling was required to distinguish between the periplasm oxidation reactions and the start of the pentose phosphate (PP) pathway, only the periplasmic secretion reactions were included in the model. The resulting fits from INCA and WUFlux indicated the conservation of the core *P. putida* metabolic features within all three strains (based on features previously reported (Kohlstedt and Wittmann, 2019; Nikel et al., 2021)), **Fig. 3, Tables S2-S4**). Namely, the ED pathway was predicted to be the exclusive catabolic route during growth on glucose, carbon reflux was observed through the cyclic ED-EMP pathway, and the pyruvate shunt was active. Within both the WT and Eng strain, the glyoxylate shunt was also inactive. The largest difference between the strains was an increase in reduced cofactor production through the periplasmic oxidized reactions and the oxidative portion of the PP pathway, helping to balance the co-factor demand imposed by indigoidine production. However, there were overall minimal flux differences between the two strains which shows that the native metabolic network accommodates indigoidine production without resulting in significant central carbon flux changes.

**Figure 3.**
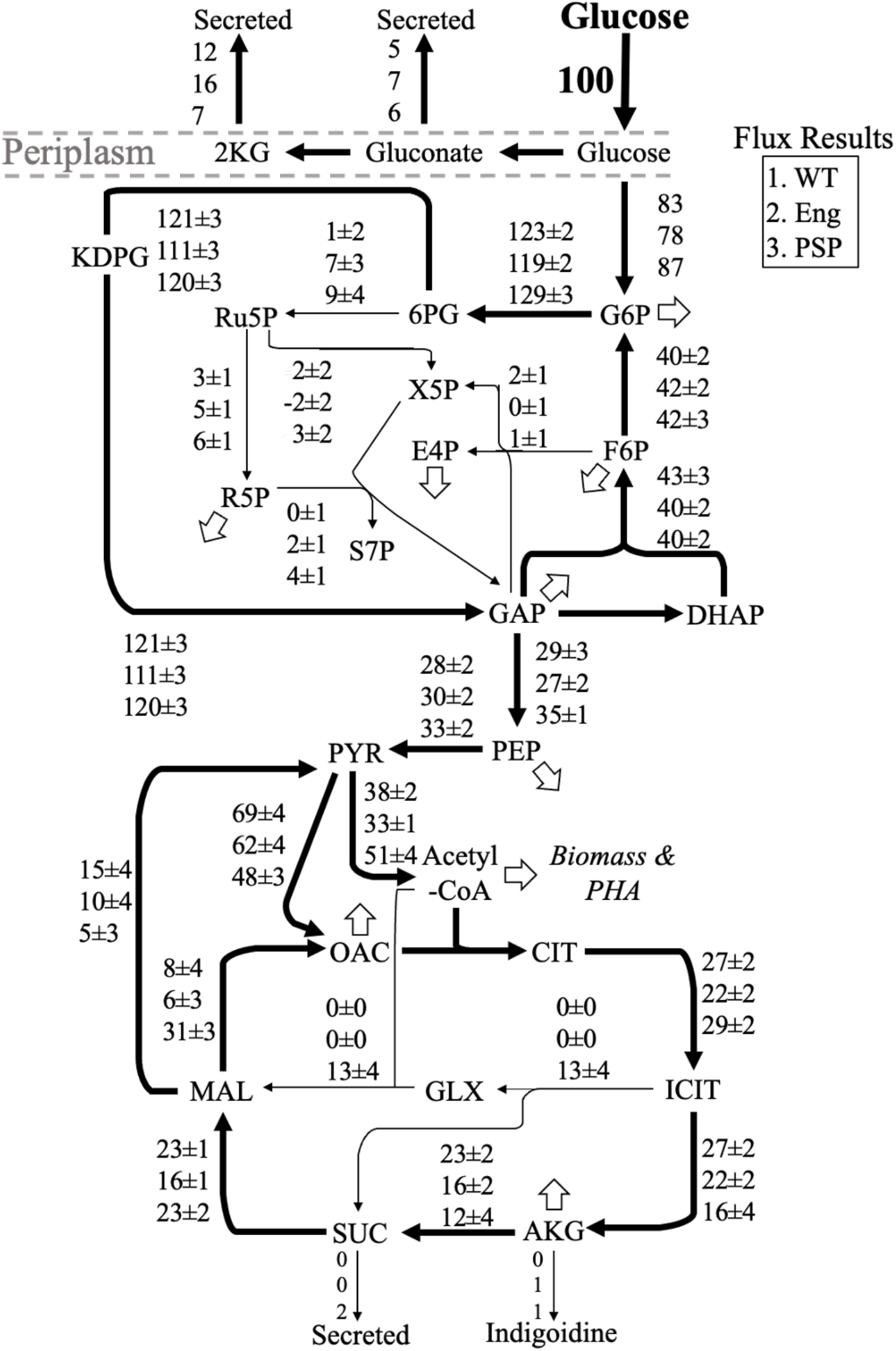
Flux networks of *P. putida* strains. Flux fitting results for the three *P. putida* strains are represented using three rows, the WT (top), Eng (middle), and PSP (bottom). Hollow arrows represent metabolite drainage for biomass equations. Some reactions are omitted from the figure for clarity. All reaction fits, uptake rates, and biomass formations can be found in the Tables S2-S4. For metabolite abbreviations, refer to the list of abbreviations within the Supplementary Material.

On the other hand, expression of the CRISPRi system impacted the flux network of the PSP strain, with flux rewiring observed in the lower half of metabolism (i.e., the TCA cycle, pyruvate shunt, **Fig. 3**). Targeting of the malic enzyme (*maeB*) achieved a 2.0–fold reduction in reaction activity of the PSP strain compared to Eng strain, with a corresponding 1.3–fold decrease in the second reaction (pyruvate carboxylase) of the pyruvate shunt. In response, 5.4–fold more flux was channeled through the malate dehydrogenase reaction (MAL → OAC). This increased activity occurred despite the targeted downregulation of three of the four *P. putida* malate dehydrogenase genes (*mdh, mqo-I, mqo-II*). The rearrangement of flux activity downstream of MAL propagated through the TCA cycle and resulted in a 1.3– to 1.4–fold reduction in flux from ICIT→AKG→SUC (**Fig. 3, Table S3-S4**) and the activation of the glyoxylate shunt. A large portion of the flux entering the TCA cycle (forming citrate) was diverted through the glyoxylate shunt pathway which potentially allowed the cell to bypass bottlenecks in the middle portion of the TCA cycle and to reduce CO_2_ carbon loss. There was a slight (1.1–fold) increase in ED glycolysis flux due to the reduction of secreted oxidized reactions, but no flux rewiring was observed in the cyclic ED-EMP pathway steps. Relatively more PP pathway flux activity in the PSP strain was also observed. However, the 95% confidence intervals overlapped for the two strains (**Table S3-S4**) and the PP pathway was predicted to have limited activity overall. Overall, these flux maps suggested that effects of multiplexed CRISPRi led to changes in the pathway activity that were dominated by a few of the down regulated targets. Next, the effects of the metabolic engineering on intracellular metabolite concentrations were investigated.

### 3.3 Intracellular metabolite concentration changes

Intracellular metabolite concentrations affect the thermodynamic driving force of enzyme reactions and buffer the flux network (Raamsdonk et al., 2001). When an enzyme abundance is altered after genetic modification, the accumulation or depletion of a metabolite can help a cell maintain its flux network. Therefore, a targeted metabolomics approach was utilized to examine the intracellular metabolite concentrations (pool sizes) in the strains. The measurements revealed a shift in pool sizes between the strains during both the growth and production phase (**Fig. 4**). In the growth phase, the Eng strain was observed to have a depletion in many central carbon metabolites compared to the WT strain. Expression of the CRISPRi knock-down construct appeared to restore metabolite pool sizes to the WT level. A significant difference (anova *p*-value < 0.05) of pool sizes were observed for erythrose 4-phosphate (E4P) with an approximate 2–fold increase in both the Eng and PSP engineered strains compared to the WT. There was also a significant increase of glyceraldehyde 3-phosphate (GAP) pool size (1.8–fold) in the PSP strain compared to both the WT and Eng strain. In the TCA cycle, the citrate (CIT) pool size of the PSP strain was increased by 1.7–fold and approximately 3.5–fold relative to the WT and Eng strains, respectively. The observed build-up of CIT in the PSP strain supports the predicted downregulation of TCA cycle enzymes and identifies the presence of a potential bottleneck in the TCA cycle. In contrast, the decrease of intracellular CIT in the Eng strain may be due to drainage of TCA cycle metabolites in order to produce indigoidine (requires two molecules of glutamine, derived from alpha-ketoglutarate, AKG). During the production phase, there was an overall reduction of most intracellular metabolites as glucose depleted. There was an approximate 3–fold increase of SUC in the PSP strain which was consistent with the observed secreted SUC. The metabolite measurements and flux observations provided insight into the robustness of the *P. putida* flux network and its capacity to accommodate both production and genetic engineering.

**Figure 4.**
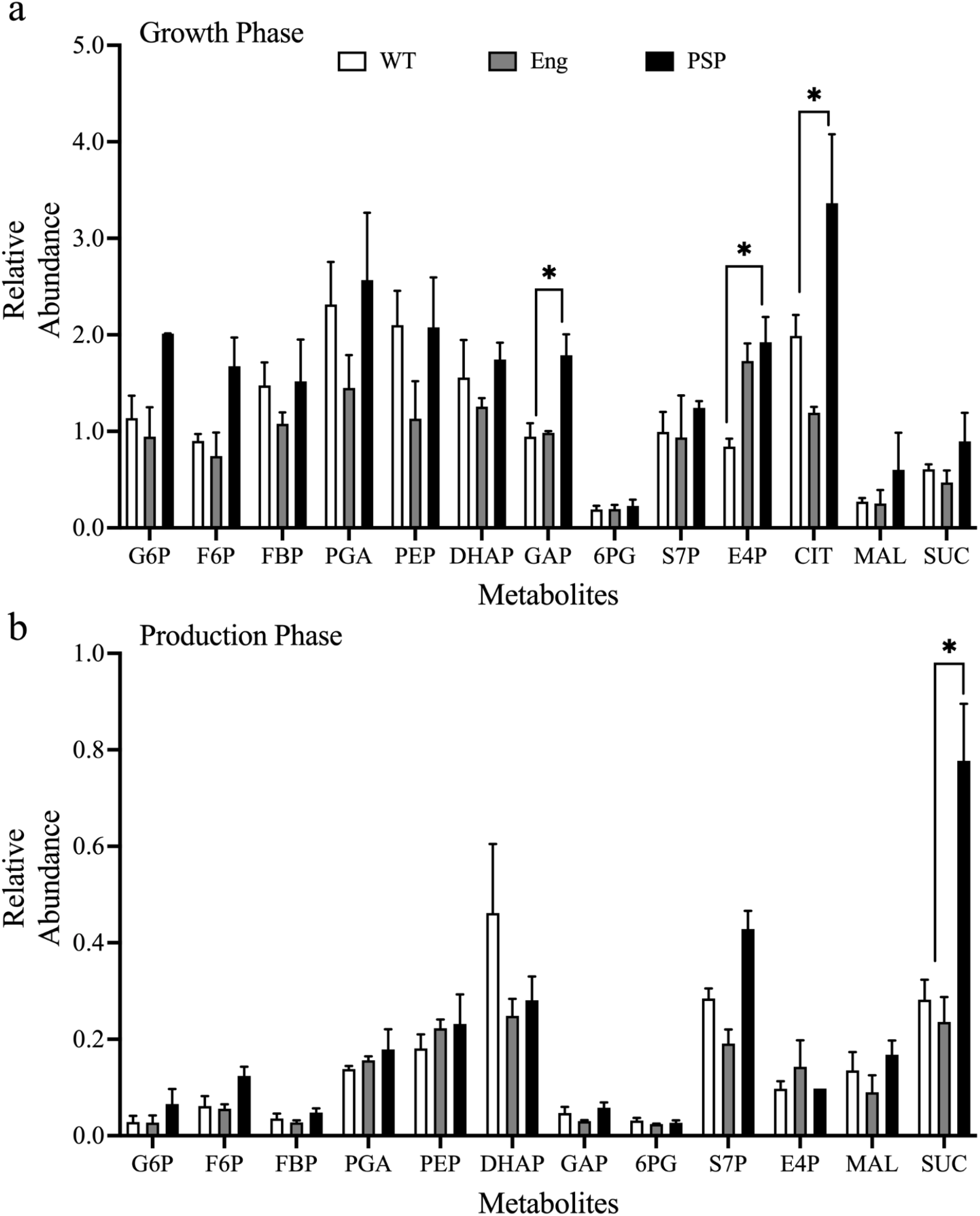
Metabolite pool sizes. Comparison of relative metabolite pool sizes in the growth (a) and production (b) phases. * denote metabolite pool sizes that were significantly different between species (p-value <0.05, one-way anova). Error bars represent the standard error (n=3), except for E4P in the PSP strain during production phase (n=1). For metabolite abbreviations, refer to the list of abbreviations within the Supplementary Information.

### 3.4 Further genetic engineering targets and strain improvements

The comprehensive omics data analysis led to the selection of four targets for further engineering. The fluxomic data suggested two major findings; first, that the glyoxylate shunt was activated and allowed carbon to bypass the central carbon metabolism indigoidine precursor, AKG and second, that downregulation of the malate dehydrogenase reaction was not sufficient despite targeting three out of four enzymes involved in the reaction by CRISPRi (*mdh, mqo-II, mqo-II*, **Fig. 3**). Thus, two genes, isocitrate dehydrogenase, *aceA* (PP_4116) and the final malate dehydrogenase, *mqo-III* (PP_0751) were selected as deletion targets. The intracellular and extracellular metabolomic data sets demonstrated a build-up and secretion of succinate from the PSP strain (**Fig. 2** and **Fig. 4**) under the specific growth conditions tested. Reutilization of the accumulated succinate would not increase indigoidine production and thus represented a loss of carbon from the system (**Fig. 2** and **Fig. S4**). The *sucC* (PP_4185) and *sucD* (PP_4186) subunits of succinyl-CoA synthetase complex were selected as the next gene targets to prevent succinate build-up and secretion. Similarly, the gluconate forming glucose dehydrogenase was deleted (ΔPP_1444) to prevent the secretion of gluconate or downstream metabolic products of gluconate (glucuronate or 3-keto-D-gulonate) during the growth phase. A final deletion strain design was included based on the previous cMCS predictions. The *phaAZC-IID* operon responsible for synthesis of the storage compound polyhydroxyalkanoate (PHA) consistently appeared as a deletion candidate in the cMCS design (Banerjee et al., 2020). The two genes targeted for downregulation (PP_5003 and PP_5005) by the CRISPRi construct catalyze the polymerization of PHA. While the PP_5003 and PP_5005 were shown to be down-regulated in the PSP strain, several other genes involved in β-oxidation (forming precursors for PHA biosynthesis) were specifically up-regulated compared to the Eng strain (Banerjee et al., 2020), **Table S5**). These genes and precursors might be sufficient to redirect carbon flux away from the indigoidine production route. Thus, the complete *pha* operon was selected for deletion.

In addition to the above deletion targets, the relatively small changes in the PSP flux network and limited growth phase production (in shaking flasks) led to the question of whether the PSP strain production could be further improved if the CRISPRi repression was more complete in its knockdown efficacy. A random mutagenesis screen of the ribosome binding site (RBS) was performed to identify a sequence that resulted in stronger expression (**see Materials and Methods 2.9**). It is well established that introducing a CRISPR system targeting a chromosomal locus for cutting leads to killing of the cells (Bikard et al., 2012). This phenotype was used to identify RBS variants for *F. novicida cpf1* expression where the concurrent introduction of a gRNA targeting *pyrF/*PP_1815 led to low CFUs in a WT strain but had no change on CFU counts in a Δ*pyrF* strain. The original RBS sequence for *cpf1* expression led to >500 CFU per μg plasmid DNA when transformed the *pyrF* locus was targeted for cutting. After screening approximately 50 RBS mutants, one candidate RBS was identified with the best cell-killing activity (~0-5 CFUs / μg plasmid DNA) implying it had improved function. The identified RBS sequence was then incorporated into the CRISPRi plasmid for glucose/indigoidine growth coupling to enhance expression (pTE442, **Table 1**). The reinforcement of the original cMCS design resulted in stronger growth coupling, with 1.6–fold (p-value=0.08) improvement in production rate during the growth-phase (6 h timepoint) compared to the PSP strain (**Fig. 5**). However, the increased production rate was not sustained over the course of cultivation and the strain only resulted in a 1.2–fold increase of titer (p-value=0.14) at the final time point (72 h).

**Figure 5.**
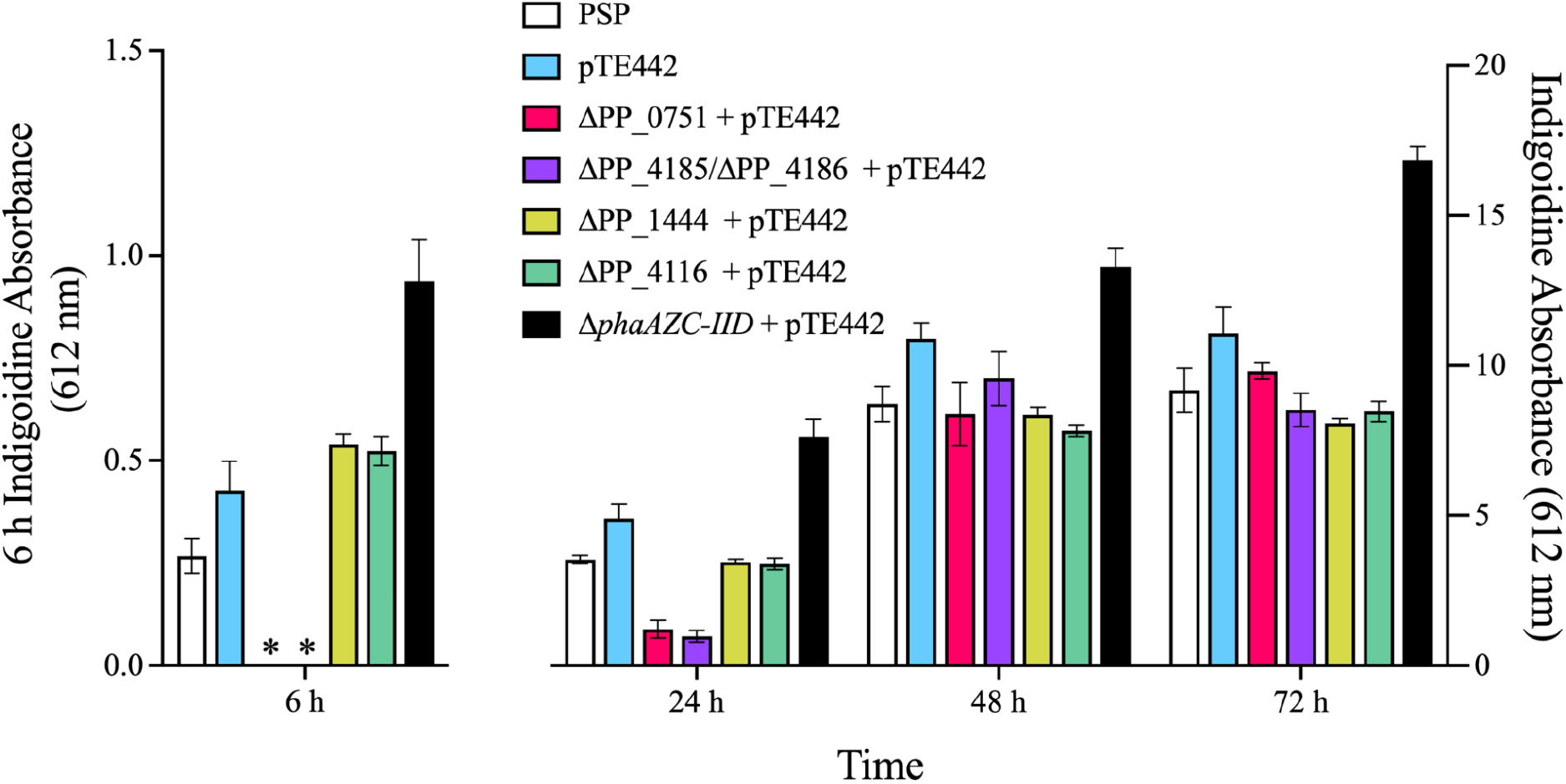
Production profiling of various indigoidine producing strains. Indigoidine titers as measured by absorbance in DMSO at 612 nm. * indicates data points that were not collected due to no observable growth. The 6 h measurements are depicted on a separate axis due to differences in the scale of the measurements. The indigoidine absorbance for the Δ*phaAZC-IID* is ~0.2 ± 0.1 g/L at 6 h and ~3.6 ± 0.2 g/L at 72 h using a previously reported standard curve (Banerjee et al., 2020). Error bars represent standard error (n ≥ 6).

Next, each of the deletion strains (ΔPP_0751, ΔPP_1444, ΔPP_4116, ΔPP_4185ΔPP_4186, Δ*phaAZC-IID*) were constructed via ssDNA recombineering (**see Materials and Methods Section 2.10**) and tested for growth-coupled indigoidine production with the RBS-optimized CRISPRi plasmid (pTE442). Deletion of the PHA operon led to a significant (p-value < 0.01) increase in both production rate and titer across the cultivation (**Fig. 5**). The Δ*phaAZC-IID* strain had a 2.2–fold increase in production during the growth phase compared to pTE442 strain, which was maintained throughout the cultivation and resulted in a 1.5–fold improvement of indigoidine titer at 72 h (p-values<0.01).

In contrast, deletion of *mqo-III* (ΔPP_0751) or the succinyl-CoA synthetase subunits (ΔPP_4185 ΔPP_4186) led to strains with substantial growth defects (reaching OD_600_ ~1–3 at 24 h) and had no improvement in production titers. RB-TnSeq analysis had previously indicated that no transposon mutants had been recovered in either of these gene loci (Eng et al., 2021; Price et al., 2018; Thompson et al., 2019). Colonies were obtained for each deletion, although in liquid media the strains exhibited a several growth defect in minimal medium. The deletion strains’ fitness defects confirmed the previous RB-TnSeq results and the study workflow aimed at avoiding potentially essential metabolic reactions targets for CRISPRi knockdown. Of the remaining deletion strains isocitrate dehydrogenase (ΔPP_4116) or glucose dehydrogenase (ΔPP_1444) did not have pronounced growth defects but had similar production metrics as the RBS-optimized growth-coupled plasmid alone at 6 h (**Fig. 5**). There was a 1.2– and 1.3–fold increase in production for the respective strains during the growth phase, albeit the increase was not significant (p-values = 0.21 and 0.17, respectively). The production rates slowed over the course of cultivation and the deletions were detrimental to overall titers with 1.3– and 1.4–fold decreases observed at 72 h (p-values <0.05). Both the ΔPP_4116 and ΔPP_1444 strains also had decreased production compared to the original PSP strain by the end of the cultivation (by 8% (p-value >0.05) and 10% (p-value <0.05), respectively). Overall, these results indicated that the modified RBS used to drive *cpf1* expression increased production 60% during the exponential phase compared to the original design while further deletion of the *pha* operon led to a 220% increase. Moreover, final product titer improvements could be detected concomitant with growth in exponential phase, in stationary phase, or both.

## 4. Discussion

### 4.1 Engineering insights on growth coupling and cellular byproducts

Growth coupling has been demonstrated as a viable strategy for improving production of non-toxic compounds like indigoidine, itaconic acid, and 1,4-butanediol (Banerjee et al., 2020; Harder et al., 2016; Yim et al., 2011). Further shifting production dynamics towards the growth phase with the improved RBS sequence continued to result in increased production and 60% more product at 6 h during the growth phase. Of the deletion strains tested, inactivating PHA synthesis was the most effective strategy and led to a 2.2–fold improvement in growth phase production and 1.5–fold more indiogine at the end of cultivation compared to the pTE442 strain (**Fig. 5**). PHAs are typically synthesized as storage compounds that are accessed under glucose starvation in *P. putida* (Ankenbauer et al., 2020) and have been previously shown to be closely tied to central carbon metabolism (de Eugenio et al., 2010b, 2010a; Escapa et al., 2012). Preventing PHA formation in *P. putida* is beneficial for indigoidine production because blocking synthesis increases acetyl-CoA and precursor flux through the TCA cycle (Escapa et al., 2012). Furthermore, PHA synthesis has also been identified as a key component in maintaining energy and redox balance by allowing for dissipation of excess energy. Disrupting this energy balancing component further drives the production of indigoidine as a means to dissipate reducing power. Overall, the *phaAZC-IID* deletions generated here prevented the accumulation of this key carbon and storage component and enabled the strain to redirect excess carbon and energy to production of indigoidine throughout the cultivation process.

However, production differences between the strains containing the improved pTE442 plasmid and the original PSP strain decreased over time (**Fig. 5**). The reduction in performance corresponded with the stoppage of biomass accumulation (**Fig. 1**), although indigoidine continued to form after glucose depletion. These observations suggest that the continued accumulation of the product after glucose exhaustion is not fully accounted for in the metabolic models and may occur as a consequence of differential gene expression (that goes beyond steady state metabolism) in the stationary phase. Extending the growth phase through a fed-batch or continuous cultivation is expected to result in further production improvements between strains. This strategy could enhance the effectiveness of the CRISPRi construct as well since many of the CRISPRi gene targets are involved in central carbon metabolism and amino acid synthesis pathways that become less active as glucose is depleted.

### 4.2 *P. putida* metabolic plasticity and flux buffering

Cellular metabolism has evolved over time to provide robust flux networks generating the precursors and energy molecules necessary for growth and survival under a variety of environmental and genetic perturbations (Czajka et al., 2020b). Changes in protein levels can be counteracted by latent pathway activation, enzyme activity changes (through post-transcriptional regulation), or metabolite levels changes (Raamsdonk et al., 2001; Wegner et al., 2015). The resulting flux buffering determines the effectiveness of CRISPRi modifications and requires integrated flux and metabolite analyses. In the PSP strain, targeting 14 genes for downregulation indirectly led to the downregulation of 167 genes and the upregulation of 139 genes compared to the Eng control strain carrying an empty vector plasmid, pTE219 (**Table 1**, **Fig. S6**). Investigating the effects of the widespread transcriptomic changes on central metabolism with ^13^C-MFA revealed enzymatic rates that were undesirable based on the original cMCS design (**Fig. 4**). Specifically, there was an increase of flux through the malate dehydrogenase reaction despite targeted downregulation of this step. It appeared that limiting flux through the malic enzyme resulted in flux rerouting through the malate dehydrogenase reaction and further perturbations downstream, indicating that the malic enzyme and the pyruvate shunt are key nodes with limited flux buffering for *P. putida* to maintain stable flux through the TCA cycle. These observations are supported by studies that showed the malic enzyme/malate dehydrogenase flux ratio shifts in response to oxidative stress (Nikel et al., 2021) or iron limitation (Sasnow et al., 2016) from its normal ratio of ~65% TCA cycle flux entering the pyruvate shunt (Kohlstedt and Wittmann, 2019; Nikel et al., 2021, 2015). A further study analyzing kinetic parameters and control coefficients in *P. putida* indicated that the malic enzyme exhibits control over the pyruvate shunt and has increased importance under stressed conditions (Tokic et al., 2020). Thus, *P. putida* appears to employ highly active anaplerotic pathways to maintain relatively stable fluxes through the core pathways.

Deletion strains generated to reduce the malate dehydrogenase activity (ΔPP_0751) and the glyoxylate shunt flux (ΔPP_4116) to match the cMCS original design resulted in a severe growth defect or reduced production at 72 h as previously discussed (**Fig. 5**). Logically extending the deletion results obtained here, it follows that implementation of the original cMCS design as a 14-gene deletion strain would have generated a strain with significantly slower growth and lower production overall compared to the multiplexed CRISPRi implementation. Considering the indigoidine production titer when PP_4116 expression is reduced *vs* its complete deletion; by RNAseq and proteomics, CRISPRi knockdown reduced PP_4116 levels by ~30% (**Table S4**). In contrast, when PP_4116 is completely abolished in the case of a gene deletion, the impact on indigoidine titer was not beneficial overall and even led to an 8% decreased production compared to the original PSP strain. While gene deletions lead to desirable re-routing fluxes in cases of simple pathways (i.e., one to two deletions), it eliminates pathways needed for the metabolic networks to accommodate further stresses. For complex engineering designs calling for multiple deletions, the resulting flux network can be too constrained and unable to maintain flexibility to compensate for both production burdens and modulation of pathways by engineered tools (Chavarría et al., 2012). Thus, utilizing CRISPRi to modulate cellular fluxes allows cells to maintain the necessary flux network flexibility that can result in stable growth and robust production in cases of complex strain designs. These CRISPRi implementations may provide an easy route to prototype strain designs but traditional full gene knockouts may be necessary to fully confirm predicted phenotypes.

## 5. Conclusions

Initial DBTL cycles can be quite effective in increasing strain production, but systems analyses are necessary to reveal cell metabolic regulations and to identify non-intuitive engineering targets in advanced cycles (Zhang et al., 2020). Here, multi-omics data provided holistic information on the *P. putida* system-wide response to complex metabolic perturbations that helped re-engineer a strain for a 50% improvement in heterologous production titer. ^13^C-MFA and the developed strains revealed the malic enzyme as a controlling node within *P. putida* for maintaining stable TCA cycle flux during stressed conditions. Broadly, multi-omic analyses can lead to understanding of metabolic regulations in light of cultivation and biomanufacturing stresses. Integrating multi-omic data into genome-scale models (Kim and Lun, 2014; Martín et al., 2015; Töpfer et al., 2015) and utilizing computational design algorithms such as cMCS (Klamt and Mahadevan, 2015) or minimization of metabolic adjustment (MOMA) (Segrè et al., 2002) can result in more robust and accurate gene targets for DBTL applications.

## Data Availability

The sequences of the plasmids generated in this study may be visualized at public-registry.jbei.org. All other data is included in the manuscript and the supplementary files.

## Author Contributions

JC DB TE YT and AM conceived the study. CY JM TE constructed and verified recombinant *P. putida* strains and plasmids. JC harvested and analyzed samples for 13C MFA analysis. NM YK collected high resolution metabolite and protein information using high resolution mass spectrometry. JC DB TE YT and AM interpreted the results. JC wrote the first draft of the manuscript and prepared figures. All authors edited and provided constructive feedback on the final manuscript. All authors have read and approved the final version of this manuscript for publication.

## Acknowledgements

We thank Victor de Lorenzo and Esteban Martínez (Centro Nacional de Biotecnología-CSIC) for insightful technical comments and sharing their recombineering plasmids and methods. JC was supported by the U.S. Department of Energy, Office of Science, Office of Workforce Development for Teachers and Scientists, Office of Science Graduate Student Research (SCGSR) program. The SCGSR program is administered by the Oak Ridge Institute for Science and Education (ORISE) for the DOE. ORISE is managed by Oak Ridge Associated Universities (ORAU) under contract number DE-SC001464. All opinions expressed in this paper are the author’s and do not necessarily reflect the policies and views of ODE, ORAU, or ORISE.

A part of this research was conducted at the Joint BioEnergy Institute (http://www.jbei.org) supported by the US Department of Energy, Office of Science, through contract DE-AC02-05CH11231 between Lawrence Berkeley National Laboratory (LBNL) and the US Department of Energy. A portion of this research was performed on a project award 10.46936/brcr.proj.2021.51792/60000322 from the Environmental Molecular Sciences Laboratory (EMSL), a DOE Office of Science User Facility sponsored by the Biological and Environmental Research program under Contract No. DE-AC05-76RL01830 at Pacific Northwest National Laboratory (PNNL). The funders had no role in study design, data collection and analysis, decision to publish, or preparation of the manuscript.

We also acknowledge the Proteomics and Mass Spectrometry Core Facility at the Donald Danforth Plant Science Center and the United States Department of Agriculture, Agricultural Research Service, for access and use of instrumentation and resources. In addition, we acknowledge support from the National Science Foundation (NSF-MCB #1616820 and NSF-DBI #1427621), the latter grant provided support for acquisition of the QTRAP LC-MS/MS used for collecting initial data in this project.

