## Supplementary Materials for "Optimizing a High Performing Multiplex-CRISPRi *P. putida* Strain with Integrated Metabolomics and ^13^C-Metabolic Flux Analyses"

### Contents

1. **List of Metabolite Abbreviations**
2. **Supplementary Figures S1-S6**
3. **Supplementary Tables S1-S5**
4. **Supplementary Methods**

### List of Metabolite Abbreviations

2KG (2KGglcn), 2-ketogluconate; 3PG, 3-phosphoglycerate; 6PG, 6-phosphoglucanate; ADP, adenosine diphosphate; Ac, acetate; AcCoA, acetyl-CoA; AKG, α-ketoglutarate; AMP, adenosine monophosphate; Ala, Alanine; Arg, Arginine; Asn, Asparagine; Asp, Asparatate; ATP, adenosine triphosphate; CIT, citrate; $CO_2$, carbon dioxide; Cys, Cysteine; DCW, dry cell weight; DHAP, dihydroxyacetone phosphate; E4P, erythrose-4-phosphate; F6P, fructose-6-phosphate; FADH2, flavin adenine dinucleotide; FBP, fructose-1,6-bisphosphate; FTHF, formyltetrahydrofolate.; FUM, fumarate; G6P, glucose-6-phosphate; GAP, glyceraldehyde-3-phosphate; Glc, glucose; Glcnt, gluconate; Gln, glutamine; Glu, glutamate; Glyox, glyoxylate; His, Histidine; ICIT, isocitrate; Ile, isoleucine; KDPG, 2-dehydro-3-deoxy-phosphogluconate; Leu, Leucine; LLDAP, l,l-diaminopimelate; Lys, Lysine; MAL, malate; Met, methionine; MEETHF, methylene-tetrahydrofolate; METHF, methyl-tetrahydrofolate; MVA, mevalonate; NADPH, nicotinamide adenine dinucleotide phosphate; NADH, nicotinamide adenine dinucleotide; P5P, pentose-5-phosphates; PEP, phosphoenol pyruvate; PGA, 3-phosphoglycerate; Phe, phenylalanine; Pro, proline; OAC, oxaloacetate; R5P, ribose-5-phosphate; Ru5P, ribulose-5-phosphate; PYR, pyruvate; S7P, sedoheptulose-7-phosphate; Ser, Serine; SUC, succinate; SucCoA, succinyl-CoA; Thr, Threonine; Tyr, Tyrosine; UDPG, uridine diphosphate glucose; UQH2, ubiquinone; Val, Valine; X5P, xylulose-5-phosphate

### Supplementary Figures

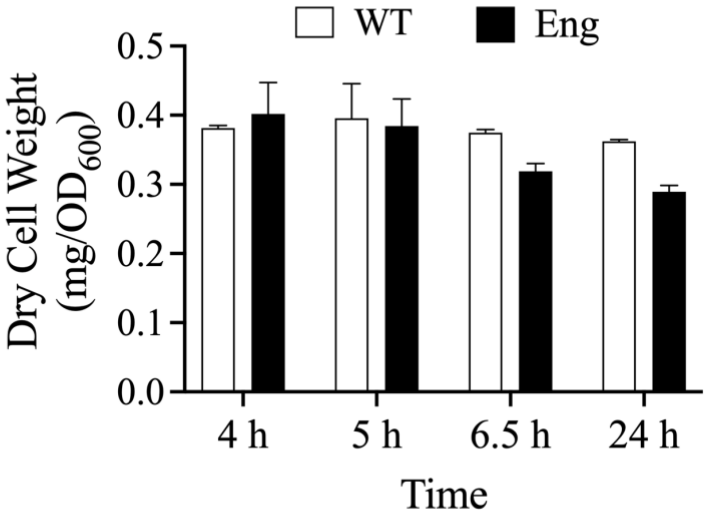


**Figure S1. Dry cell weight per $OD_{600}$ measured over the course of growth.** The DCW was measured in the Engineered and indigoidine producing strains grown in 50 mL cultures. The biomass was dried by lyophilization. The data was used to calculate intracellular metabolite concentrations per unit biomass. Error bars represent standard error (n=2).

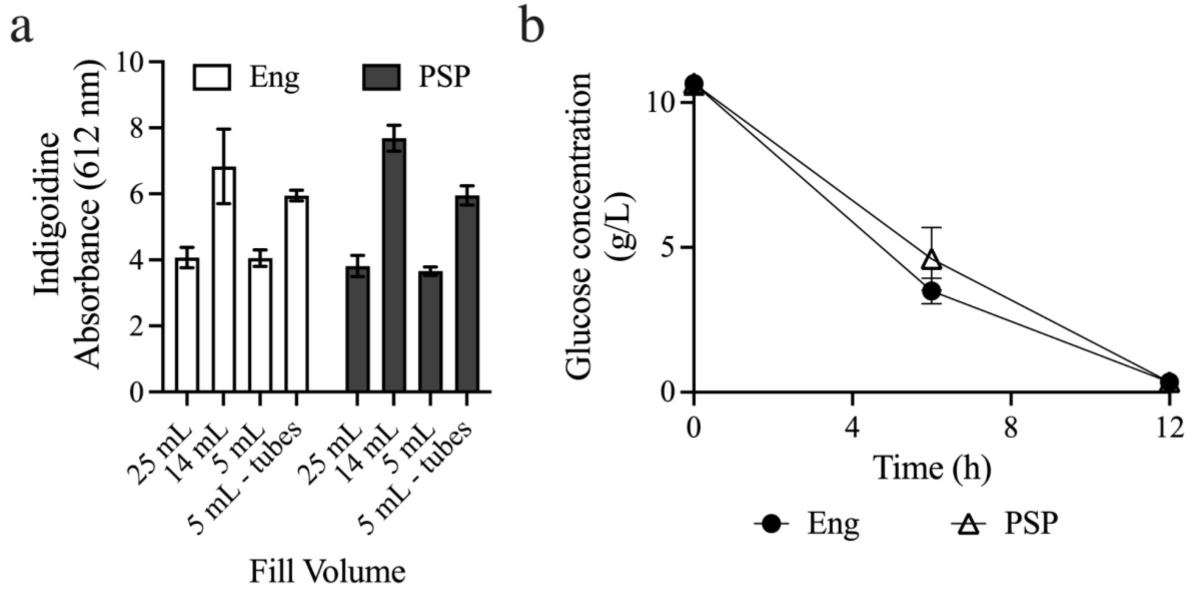


**Figure S2. Strain production verification. (a)** Screening across a variety of scales to locate the best format for flux analysis. **(b)** Glucose consumption curves for the Eng strain (6.6 ± 0.4 mmol/h) and PSP strain (5.6 ± 1.0 mmol/h). Error bars represent the standard error (a, n=4; b, n=3).

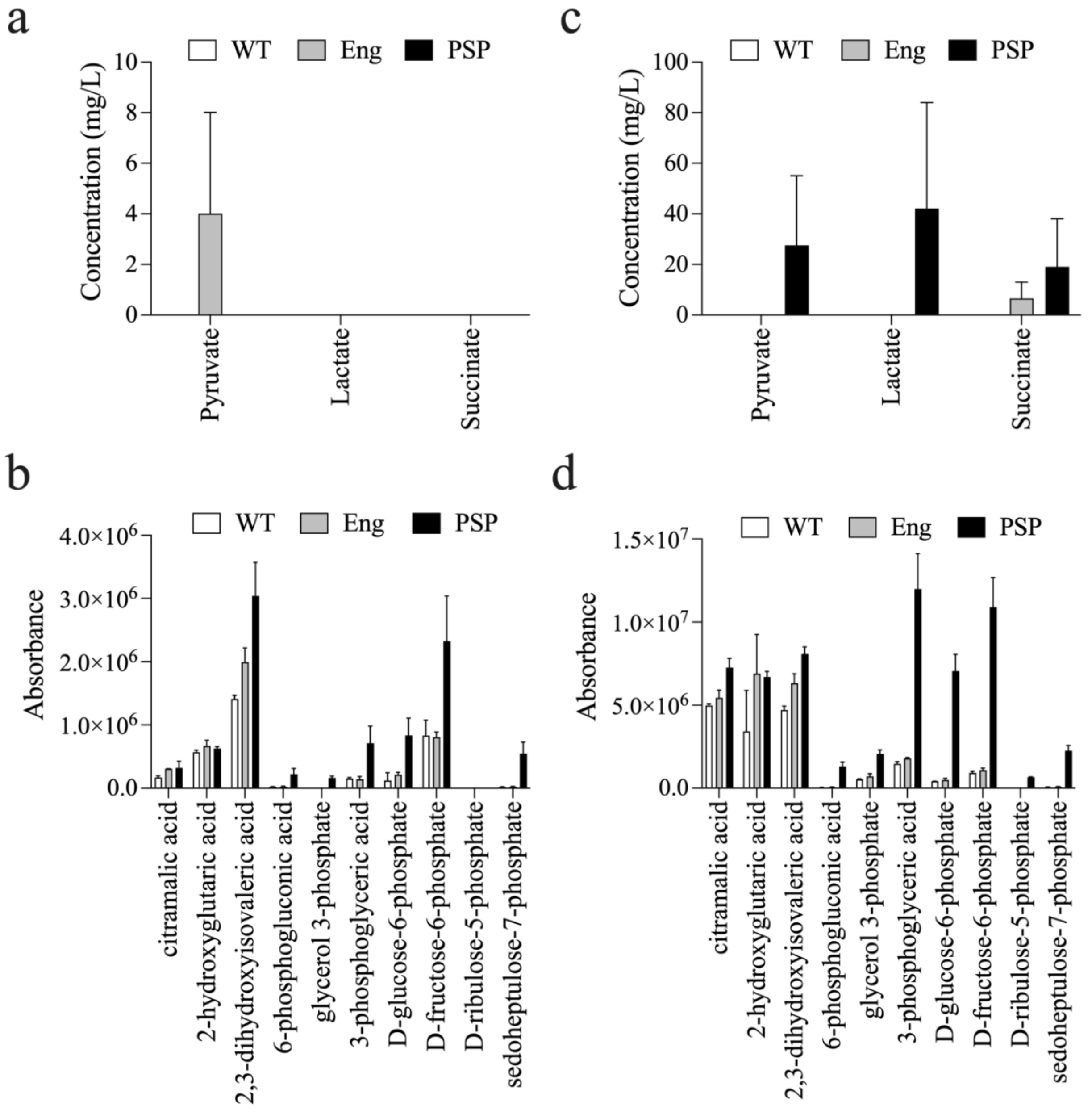


**Figure S3. Detected Extracellular metabolites. a and b.** Growth phase detected metabolites. **c and d.** Production phase detected metabolites. From the figure above, it was observed that the PSP had unique organic acid secretion at the production phase measurement (b). Absorbance cannot be used to determine absolute concentrations. Sugar phosphate presences are likely due to metabolite leakage. Error bars represent standard error (n=2).

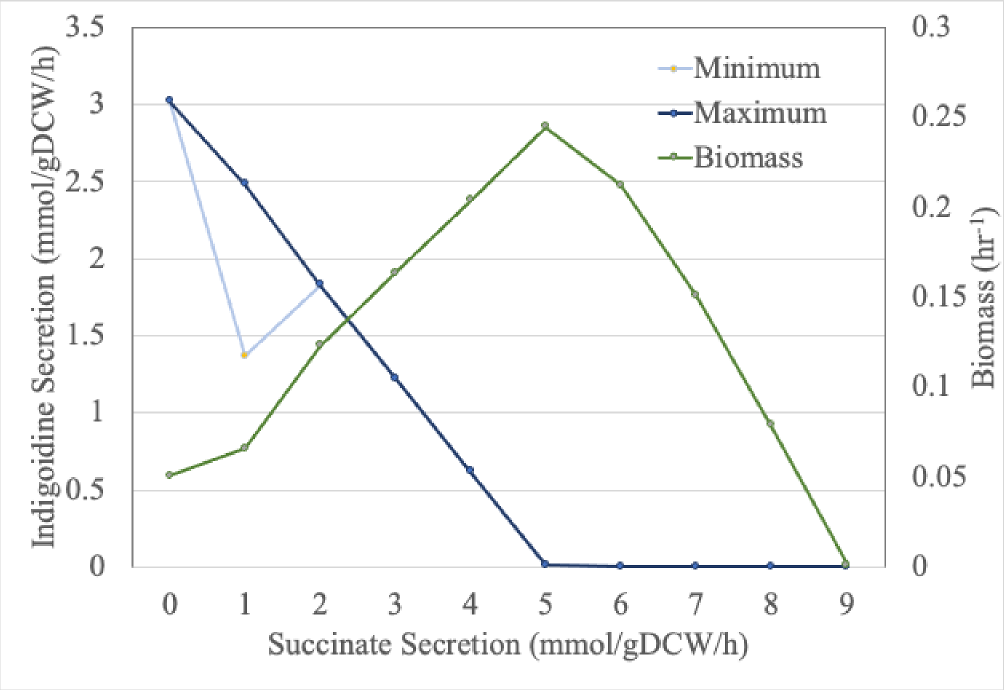


**Figure S4 Effect of Succinate secretion on the cMCS based original PSP design.** A robustness analysis followed by flux variability analysis (FVA) was performed to simulate the effect of succinate secretion on biomass as well as product substrate pairing for indigoidine production in the PSP design.

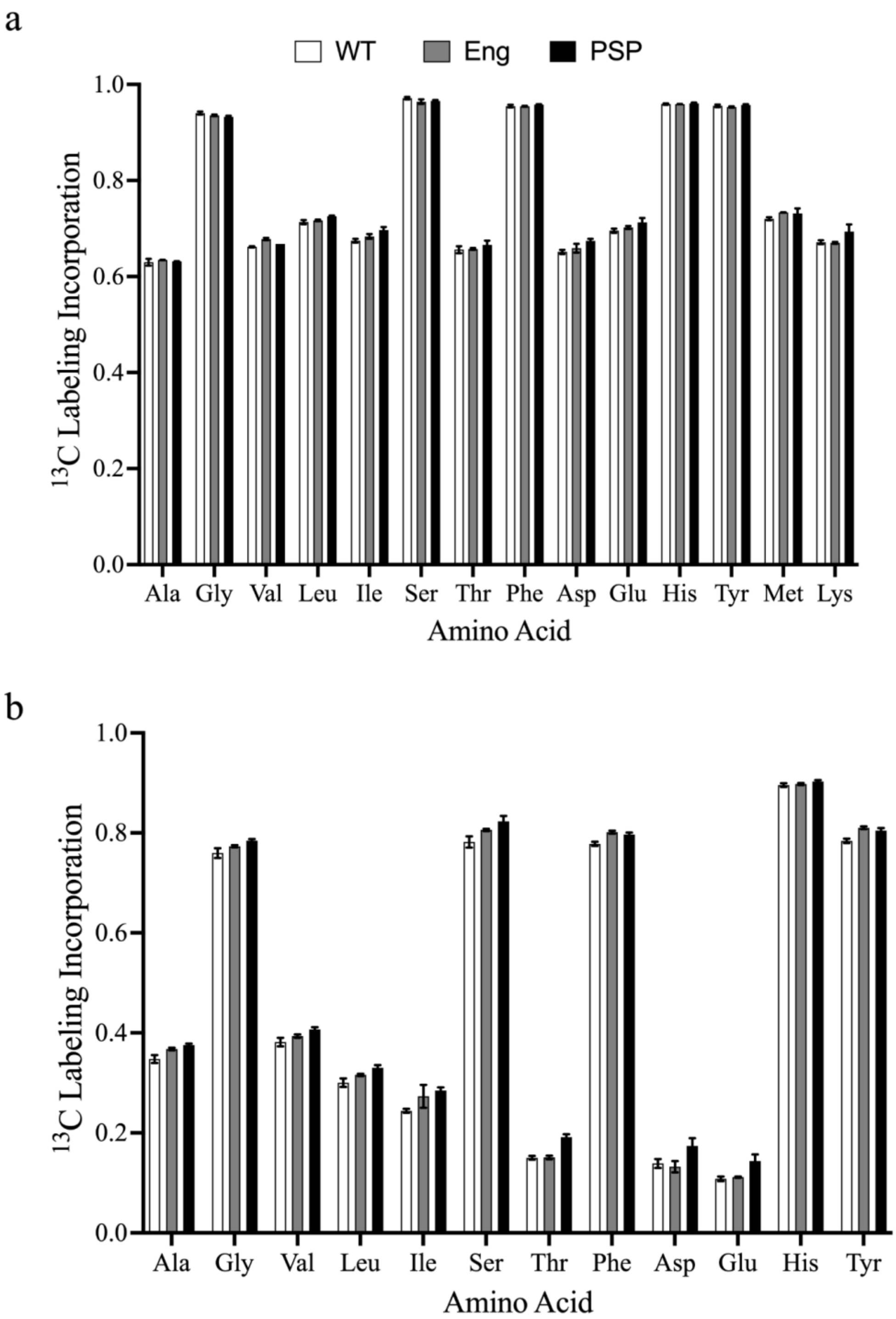


**Figure S5. Amino acid total label for the fragments containing each carbon from the amino acids. (a**) labeling experiment one conducted with one labeled carbon source (1,2-$^{13}C_2$ glucose). **(b)** labeling experiment two conducted with a mixture of two labeled carbon sources (1,2-$^{13}C_2$ glucose, U-$^{13}C_6$ glucose). Error bars represent the standard error (A; n=2, B; n=3).

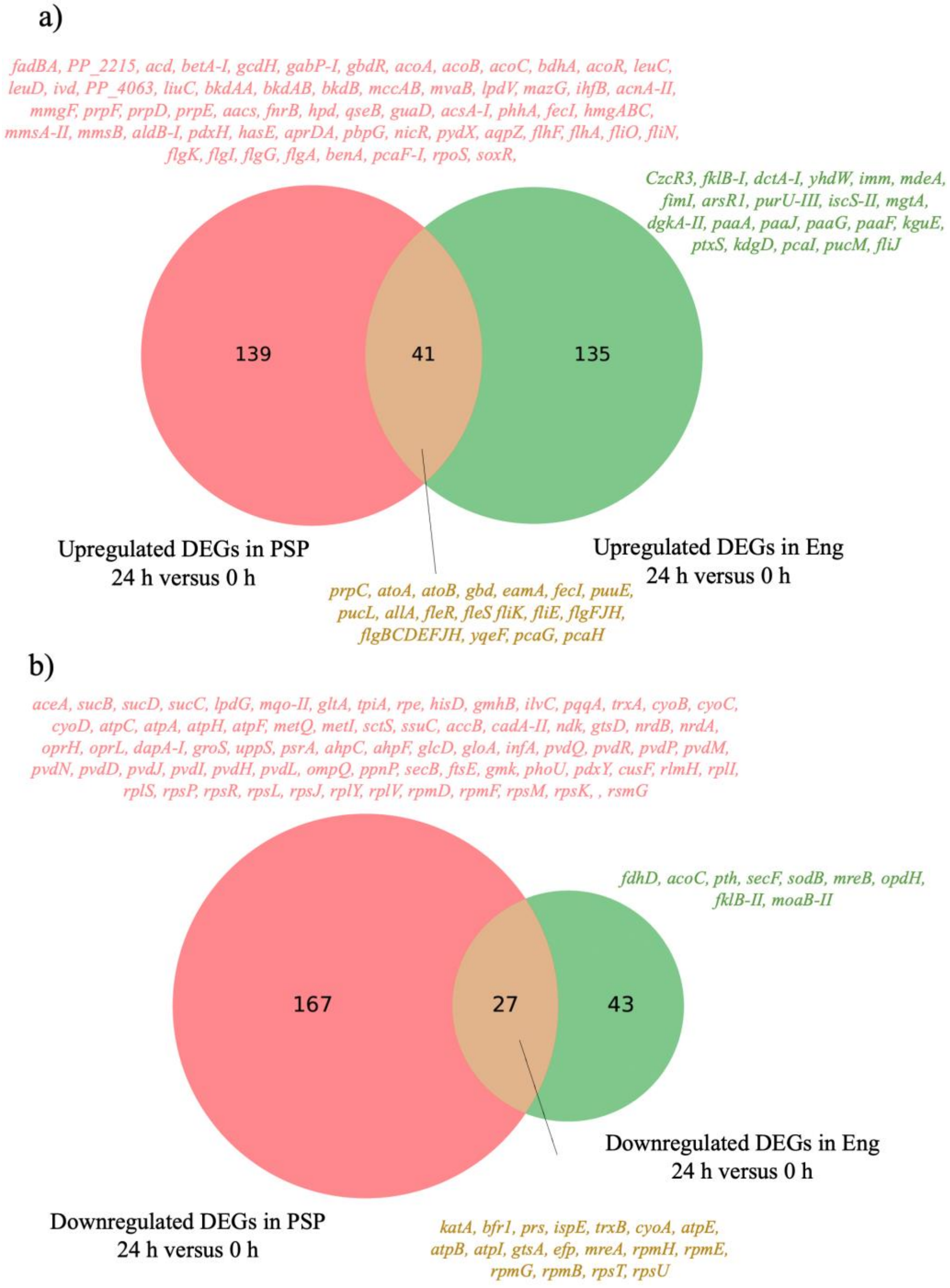


**Figure S6. Differentially Expressed Genes (DEGs) for the PSP strain compared to Eng strain.** First, DEGs were calculated using RNA-sequencing data; Log2 ratios of 24 h divided by 0 hr for PSP and Eng, respectively. Then, upregulated and downregulated genes were identified based on a fold change cutoff of 4 (|log2| >2) and a *p*-value < 0.001. The above venn diagrams show common (brown) and unique genes for PSP strain (Pink) and Eng (Green) strain.

### Supplementary Tables

**Table S1. Oligos used in this study.**

| Primer Number | Sequence (5' - 3') | Notes |
|---|---|---|
| TEAM-2260 | CATCCCGTATGAGTGTCTGAACC | PP_0751 colony PCR |
| TEAM-2261 | GCGCCGCACCTGCGGGCATTACCGA | PP_0751 colony PCR |
| TEAM-2187 | TATGCTGTCAATTAGGTAGCTGTAGATTGTGGAGACGCGTTAAGCCGGTCTGAAACGCAAAAACGCCCCTCGGGGCGTTTTTTGTGCCT | PP_0751 recombineering oligo, deletion of PP_0751 |
| TEAM-2144 | GAGCAGTTTCTAACCCCTTTTTTGACCTGACGGAGAACCTACGGCGACACCGCTCCCGCAGGCTCAACCCTGAGGGAGCGGCCTTGTGT | PP_1444 recombineering oligo, deletion of PP_1444 |
| TEAM-2158 | CCGACTGGCTCACGGTACGCCC | PP_1444 colony PCR primer |
| TEAM-2159 | GAGCAGCCTTGTGTCGCGATTGGGC | PP_1444 colony PCR primer |
| TEAM-2619 | CGCCACAGCAACCGGTACTCGTCTCAGGACAACGGAGCGTCGTAGGCCTGCTGGAGATGTAGTGTTGCAGCCGACGCATTCGCGGGTAAA | recombineering oligo to delete phaAZC-*II*D operon |
| TEAM-2621 | CTGGCACCGGTACCCTGATCTGA | PP_5005 / phaAZC-IID colony PCR primer |
| TEAM-2622 | CAACCCTTGGGCCTGTGGTGAT | PP_5005 / phaAZC-IID colony PCR primer |
| TEAM-2626 | TGGCGCAGCGCCGAAGCGCAGTACCTAACGAAGACGGTAAAAAGCTACTGCTACACCCCCCACGCGCACAGAGGCCACCTTCGGGTGGCC | recombineering oligo to delete PP_4185 PP_4186 succinate synthetase complex |
| TEAM-2627 | GAAAGTCCGCCGGACCGGATC | PP_4186 colony PCR primer |
| TEAM-2628 | AATCTCTATTGCCAGCACCGGCC | PP_4186 colony PCR primer |

| | | |
|---|---|---|
| TEAM-2634 | CATCAGCACAATTAACCCAAGATCGATGCGACGAGGTTTATTTCCTAGTGGGTTGCCGTTGAACAAAGGCCCGGGGCTTGTGAAAGCTCC | recombineering oligo to delete PP_4116 isocitrate lyase succinate synthetase complex |
| TEAM-2631 | ATTCCTGCGAACAGGACAGCCG | PP_4116 colony PCR primer |
| TEAM-2632 | CTCCCGATCAGTGTAGGTGCGG | PP_4116 colony PCR primer |

**Table S2. INCA WT Strain Flux Fitting**

| Reaction | Best Fit | Std Err | LB | UB |
|---|---|---|---|---|
| Glc.ex -> Glc.pp | 100 | 0 | 100 | 100 |
| Glc.pp -> Glcnt.pp + UQH2 | 17 | 0 | 17 | 17 |
| Glcnt.pp -> 2KGglcn.pp + FADH2 | 12 | 0 | 12 | 12 |
| Glc.pp + 2*ATP -> G6P | 83 | 0 | 83 | 83 |
| Glcnt.pp -> Glcnt.ex | 5 | 0 | 5 | 5 |
| 2KGglcn.pp -> 2KGglcn.ex | 12 | 0 | 12 | 12 |
| G6P -> 6PG + NADPH | 123 | 2 | 119 | 127 |
| 6PG -> KDPG | 121 | 3 | 118 | 126 |
| KDPG -> Pyr + GAP | 121 | 3 | 118 | 126 |
| 6PG -> Ru5P + CO2 + NADPH | 1 | 2 | 0 | 6 |

| | | | | |
|---|---|---|---|---|
| Ru5P <-> X5P | -2 | 2 | -3 | 1 |
| Ru5P <-> R5P | 3 | 1 | 3 | 5 |
| R5P + X5P <-> S7P + GAP | 0 | 1 | -1 | 2 |
| E4P + X5P <-> F6P + GAP | -2 | 1 | -3 | -1 |
| S7P + GAP <-> E4P + F6P | 0 | 1 | -1 | 2 |
| G6P <-> F6P | -40 | 2 | -45 | -37 |
| FBP -> F6P | 43 | 3 | 39 | 47 |
| FBP <-> DHAP + GAP | -43 | 3 | -47 | -39 |
| GAP <-> DHAP | 43 | 3 | 39 | 47 |
| GAP <-> 3PG + ATP + NADH | 34 | 2 | 29 | 37 |
| 3PG <-> PEP | 29 | 3 | 24 | 32 |
| PEP -> Pyr + ATP | 28 | 2 | 24 | 32 |
| Pyr + ATP -> PEP | 5 | 2 | 3 | 7 |
| Pyr -> AcCoA + CO2 + NADH | 38 | 2 | 36 | 45 |
| OAC + AcCoA <-> Cit | 27 | 2 | 25 | 29 |
| Cit <-> ICit | 27 | 2 | 25 | 29 |
| ICit -> AKG + CO2 + NADPH | 27 | 2 | 20 | 28 |
| AKG -> SucCoA + CO2 + NADH | 23 | 2 | 15 | 24 |

| | | | | |
|---|---|---|---|---|
| SucCoA -> Suc + ATP | 23 | 2 | 8 | 24 |
| Suc -> Fum + FADH2 | 23 | 2 | 19 | 24 |
| Fum <-> Mal | 23 | 1 | 20 | 24 |
| Mal -> OAC + NADH | 0 | 0 | 0 | 16 |
| Mal -> OAC + FADH2 | 8 | 4 | 0 | 19 |
| ICit -> Glyox + Suc | 0 | 0 | 0 | 8 |
| Glyox + AcCoA -> Mal | 0 | 0 | 0 | 8 |
| PEP + CO2 -> OAC | 0 | 2 | 0 | 3 |
| Pyr + CO2 + ATP -> OAC | 69 | 4 | 57 | 77 |
| Mal -> Pyr + CO2 + NADPH | 15 | 4 | 8 | 22 |
| AKG + NADPH + NH3 -> Glu | 89 | 3 | 87 | 93 |
| Glu + ATP + NH3 -> Gln | 0 | 2 | NaN | 4 |
| Glu + ATP + 2*NADPH -> Pro | 0 | 0 | 0 | 3 |
| Glu + CO2 + Gln + Asp + AcCoA + 5*ATP + NADPH -> Arg + AKG + Fum + Ac | 0 | 0 | 0 | 4 |
| OAC + Glu -> Asp + AKG | 44 | 1 | 41 | 49 |
| Asp + 2*ATP + NH3 -> Asn | 0 | 0 | 0 | 9 |
| Pyr + Glu -> Ala + AKG | 0 | 0 | 0 | 8 |
| 3PG + Glu -> Ser + AKG + NADH | 0 | 2 | 0 | 2 |

| | | | | |
|---|---|---|---|---|
| Ser <-> Gly + MEETHF | 0 | 2 | 0 | 2 |
| Gly <-> CO2 + MEETHF + NADH + NH3 | 0 | 0 | 0 | 0 |
| Thr <-> Gly + AcCoA + NADH | 0 | 2 | -2 | 0 |
| Ser + AcCoA + 3*ATP + 4*NADPH + SO4 -> Cys + Ac | 0 | 0 | 0 | 1 |
| Asp + Pyr + Glu + SucCoA + ATP + 2*NADPH -> LLDAP + AKG + Suc | 0 | 0 | 0 | 12 |
| LLDAP -> Lys + CO2 | 0 | 0 | 0 | 12 |
| Asp + 2*ATP + 2*NADPH -> Thr | 44 | 2 | 33 | 45 |
| Asp + METHF + Cys + SucCoA + ATP + 2*NADPH -> Met + Pyr + Suc_int + NH3 | 0 | 0 | 0 | 1 |
| Pyr + Pyr + Glu + NADPH -> Val + CO2 + AKG | 0 | 0 | 0 | 4 |
| AcCoA + Pyr + Pyr + Glu + NADPH -> Leu + CO2 + CO2 + AKG + NADH | 0 | 0 | 0 | 1 |
| Thr + Pyr + Glu + NADPH -> Ile + CO2 + AKG + NH3 | 44 | 1 | 33 | 45 |
| PEP + PEP + E4P + Glu + ATP + NADPH -> Phe + CO2 + AKG | 1 | 1 | 0 | 2 |
| PEP + PEP + E4P + Glu + ATP + NADPH -> Tyr + CO2 + AKG + NADH | 0 | 0 | 0 | 2 |
| Ser + R5P + PEP + E4P + PEP + Gln + 3*ATP + NADPH -> Trp + CO2 + GAP + Pyr + Glu | 0 | 0 | 0 | 1 |
| R5P + FTHF + Gln + Asp + 5*ATP -> His + AKG + Fum + 2*NADH | 0 | 2 | 0 | 2 |
| MEETHF + NADH -> METHF | 0 | 0 | 0 | 1 |
| MEETHF -> FTHF + NADPH | 0 | 2 | 0 | 2 |

| | | | | |
|---|---|---|---|---|
| NADH <-> NADPH | 58 | 4 | 47 | 68 |
| NH3.ext -> NH3 | 45 | 4 | 44 | 59 |
| SO4.ext -> SO4 | 0 | 0 | 0 | 1 |
| O2.ext -> O2 | 96 | 2 | 90 | 102 |
| CO2_unlabeled <-> CO2 | -80 | 1426 | -82 | 67 |
| CO2 -> CO2.ex | 0 | 1426 | 0 | 13 |
| ATP -> ATP.maintenance | 1.0 | 0.0 | 0.9 | 1.0 |
| NADPH -> NADPH.maintenance | 0.7 | 0.0 | 0.6 | 0.7 |
| NADH + O2 -> 3*ATP | 37 | 4 | 28 | 49 |
| FADH2 + O2 -> 2*ATP | 42 | 4 | 31 | 53 |
| UQH2 + O2 -> 3*ATP | 17 | 0 | 17 | 17 |
| 1.42*G6P + 0.56*F6P + 5.47*R5P + 2.77*E4P + 0.88*GAP + 8.86*3PG + 5.84*PEP + 14.35*Pyr + 20.49*AcCoA + 7.83*AKG + 11.04*OAC -> Biomass | 0.55 | 0 | 0.55 | 0.55 |
| SSR | 158 | | 113 | 179 |

#### Table S3. INCA Eng Strain Flux Fitting

| Reaction | Best Fit | Std Err | LB | UB |
|---|---|---|---|---|
| Glc.ex -> Glc.pp (6.6 ± 0.4 mmol/h) | 100 | 0 | 100 | 100 |
| Glc.pp -> Glcnt.pp + UQH2 | 23 | 0 | 23 | 23 |
| Glcnt.pp -> 2KGglcn.pp + FADH2 | 16 | 0 | 16 | 16 |
| Glc.pp + 2*ATP -> G6P | 78 | 0 | 78 | 78 |
| Glcnt.pp -> Glcnt.ex | 7 | 0 | 7 | 7 |
| 2KGglcn.pp -> 2KGglcn.ex | 16 | 0 | 16 | 16 |
| G6P -> 6PG + NADPH | 119 | 2 | 112 | 123 |
| 6PG -> KDPG | 111 | 3 | 106 | 116 |
| KDPG -> Pyr + GAP | 111 | 3 | 106 | 116 |
| 6PG -> Ru5P + CO2 + NADPH | 7 | 3 | 0 | 12 |
| Ru5P <-> X5P | 2 | 2 | -2 | 5 |
| Ru5P <-> R5P | 5 | 1 | 3 | 7 |
| R5P + X5P <-> S7P + GAP | 2 | 1 | 0 | 4 |
| E4P + X5P <-> F6P + GAP | 0 | 1 | -2 | 2 |

| | | | | |
|---|---|---|---|---|
| S7P + GAP <-> E4P + F6P | 2 | 1 | 0 | 4 |
| G6P <-> F6P | -42 | 2 | -46 | -36 |
| FBP -> F6P | 40 | 2 | 35 | 44 |
| FBP <-> DHAP + GAP | -40 | 2 | -44 | -35 |
| GAP <-> DHAP | 40 | 2 | 35 | 44 |
| GAP <-> 3PG + ATP + NADH | 31 | 2 | 27 | 35 |
| 3PG <-> PEP | 27 | 2 | 22 | 31 |
| PEP -> Pyr + ATP | 30 | 2 | 26 | 34 |
| Pyr + ATP -> PEP | 8 | 1 | 5 | 10 |
| Pyr -> AcCoA + CO2 + NADH | 33 | 1 | 31 | 45 |
| OAC + AcCoA <-> Cit | 22 | 2 | 19 | 24 |
| Cit <-> ICit | 22 | 2 | 19 | 24 |
| ICit -> AKG + CO2 + NADPH | 22 | 2 | 11 | 23 |
| AKG -> SucCoA + CO2 + NADH | 16 | 2 | 4 | 18 |
| SucCoA -> Suc + ATP | 16 | 2 | 0 | 17 |
| Suc -> Fum + FADH2 | 16 | 2 | 12 | 18 |
| Fum <-> Mal | 16 | 1 | 13 | 18 |
| Mal -> OAC + NADH | 0 | 0 | 0 | 19 |

| | | | | |
|---|---|---|---|---|
| Mal -> OAC + FADH2 | 6 | 3 | 0 | 20 |
| ICit -> Glyox + Suc | 0 | 0 | 0 | 10 |
| Glyox + AcCoA -> Mal | 0 | 0 | 0 | 10 |
| PEP + CO2 -> OAC | 1 | 2 | 0 | 4 |
| Pyr + CO2 + ATP -> OAC | 62 | 4 | 46 | 69 |
| Mal -> Pyr + CO2 + NADPH | 10 | 4 | 1 | 18 |
| AKG + NADPH + NH3 -> Glu | 83 | 1 | 80 | 86 |
| Glu + ATP + NH3 -> Gln | 2 | 1 | NaN | 6 |
| Glu + ATP + 2*NADPH -> Pro | 0 | 0 | 0 | 3 |
| Glu + CO2 + Gln + Asp + AcCoA + 5*ATP + NADPH -> Arg + AKG + Fum + Ac | 0 | 0 | 0 | 4 |
| OAC + Glu -> Asp + AKG | 41 | 1 | 36 | 46 |
| Asp + 2*ATP + NH3 -> Asn | 0 | 0 | 0 | 9 |
| Pyr + Glu -> Ala + AKG | 0 | 0 | 0 | 6 |
| 3PG + Glu -> Ser + AKG + NADH | 0 | 1 | 0 | 2 |
| Ser <-> Gly + MEETHF | 0 | 1 | 0 | 1 |
| Gly <-> CO2 + MEETHF + NADH + NH3 | 0 | 0 | 0 | 0 |
| Thr <-> Gly + AcCoA + NADH | 0 | 1 | -2 | 0 |
| Ser + AcCoA + 3*ATP + 4*NADPH + SO4 -> Cys + Ac | 0 | 0 | 0 | 1 |

| | | | | |
|---|---|---|---|---|
| Asp + Pyr + Glu + SucCoA + ATP + 2*NADPH -> LLDAP + AKG + Suc | 0 | 0 | 0 | 11 |
| LLDAP -> Lys + CO2 | 0 | 0 | 0 | 11 |
| Asp + 2*ATP + 2*NADPH -> Thr | 41 | 1 | 29 | 41 |
| Asp + METHF + Cys + SucCoA + ATP + 2*NADPH -> Met + Pyr + Suc_int + NH3 | 0 | 0 | 0 | 1 |
| Pyr + Pyr + Glu + NADPH -> Val + CO2 + AKG | 0 | 0 | 0 | 3 |
| AcCoA + Pyr + Pyr + Glu + NADPH -> Leu + CO2 + CO2 + AKG + NADH | 0 | 0 | 0 | 1 |
| Thr + Pyr + Glu + NADPH -> Ile + CO2 + AKG + NH3 | 41 | 1 | 31 | 41 |
| PEP + PEP + E4P + Glu + ATP + NADPH -> Phe + CO2 + AKG | 0 | 0 | 0 | 2 |
| PEP + PEP + E4P + Glu + ATP + NADPH -> Tyr + CO2 + AKG + NADH | 0 | 1 | 0 | 2 |
| Ser + R5P + PEP + E4P + PEP + Gln + 3*ATP + NADPH -> Trp + CO2 + GAP + Pyr + Glu | 0 | 0 | 0 | 1 |
| R5P + FTHF + Gln + Asp + 5*ATP -> His + AKG + Fum + 2*NADH | 0 | 1 | 0 | 1 |
| MEETHF + NADH -> METHF | 0 | 0 | 0 | 1 |
| MEETHF -> FTHF + NADPH | 0 | 1 | 0 | 1 |
| NADH <-> NADPH | 48 | 5 | 37 | 63 |
| NH3.ext -> NH3 | 45 | 2 | 44 | 57 |
| SO4.ext -> SO4 | 0 | 0 | 0 | 1 |
| O2.ext -> O2 | 90 | 2 | 83 | 95 |

| | | | | |
|---|---|---|---|---|
| CO2_unlabeled <-> CO2 | -66 | 185 | -67 | 42 |
| CO2 -> CO2.ex | 0 | 185 | 0 | 133 |
| ATP -> ATP.maintenance | 1 | 0 | 1 | 1 |
| NADPH -> NADPH.maintenance | 1 | 0 | 1 | 1 |
| NADH + O2 -> 3*ATP | 32 | 4 | 20 | 43 |
| FADH2 + O2 -> 2*ATP | 35 | 4 | 26 | 48 |
| UQH2 + O2 -> 3*ATP | 23 | 0 | 23 | 23 |
| 2*Gln + 2*ATP + 2*FADH2 -> Indigo | 1 | 0 | 1 | 1 |
| 1.42*G6P + 0.56*F6P + 5.47*R5P + 2.77*E4P + 0.88*GAP + 8.86*3PG + 5.84*PEP + 14.35*Pyr + 20.49*AcCoA + 7.83*AKG + 11.04*OAC -> Biomass | 0.55 | 0.00 | 0.55 | 0.55 |
| SSR | 158 | | 105 | 169 |

**Table S3. INCA PSP Strain Flux Fitting**

| Reaction | Best Fit | Std Err | LB | UB |
|---|---|---|---|---|
| Glc.ex -> Glc.pp (5.6 ± 1.0 mmol/h) | 100 | 0 | 100 | 100 |
| Glc.pp -> Glcnt.pp + UQH2 | 13 | 0 | 13 | 13 |
| Glcnt.pp -> 2KGglcn.pp + FADH2 | 7 | 0 | 7 | 7 |
| Glc.pp + 2*ATP -> G6P | 87 | 0 | 87 | 87 |
| Glcnt.pp -> Glcnt.ex | 6 | 0 | 6 | 6 |
| 2KGglcn.pp -> 2KGglcn.ex | 7 | 0 | 7 | 7 |
| G6P -> 6PG + NADPH | 129 | 3 | 124 | 134 |
| 6PG -> KDPG | 120 | 3 | 116 | 129 |
| KDPG -> Pyr + GAP | 120 | 3 | 116 | 129 |
| 6PG -> Ru5P + CO2 + NADPH | 9 | 4 | 0 | 13 |
| Ru5P <-> X5P | 3 | 2 | -4 | 5 |
| Ru5P <-> R5P | 6 | 1 | 2 | 7 |
| R5P + X5P <-> S7P + GAP | 4 | 1 | 1 | 5 |
| E4P + X5P <-> F6P + GAP | -1 | 1 | -3 | 1 |
| S7P + GAP <-> E4P + F6P | 4 | 1 | 1 | 5 |

| | | | | |
|---|---|---|---|---|
| G6P <-> F6P | -42 | 3 | -48 | -38 |
| FBP -> F6P | 40 | 2 | 37 | 45 |
| FBP <-> DHAP + GAP | -40 | 2 | -45 | -37 |
| GAP <-> DHAP | 40 | 2 | 37 | 45 |
| GAP <-> 3PG + ATP + NADH | 39 | 1 | 35 | 42 |
| 3PG <-> PEP | 35 | 1 | 30 | 38 |
| PEP -> Pyr + ATP | 33 | 2 | 29 | 37 |
| Pyr + ATP -> PEP | 7 | 1 | 4 | 10 |
| Pyr -> AcCoA + CO2 + NADH | 51 | 4 | 44 | 60 |
| OAC + AcCoA <-> Cit | 29 | 2 | 25 | 33 |
| Cit <-> ICit | 29 | 2 | 25 | 33 |
| ICit -> AKG + CO2 + NADPH | 16 | 4 | 9 | 26 |
| AKG -> SucCoA + CO2 + NADH | 12 | 4 | 4 | 19 |
| SucCoA -> Suc + ATP | 12 | 4 | 0 | 20 |
| Suc -> Fum + FADH2 | 23 | 2 | 19 | 28 |
| Fum <-> Mal | 23 | 2 | 19 | 27 |
| Mal -> OAC + NADH | 0 | 0 | 0 | 34 |
| Mal -> OAC + FADH2 | 31 | 3 | 0 | 37 |

| | | | | |
|---|---|---|---|---|
| ICit -> Glyox + Suc | 13 | 4 | 6 | 20 |
| Glyox + AcCoA -> Mal | 13 | 4 | 6 | 20 |
| PEP + CO2 -> OAC | 0 | 2 | 0 | 3 |
| Pyr + CO2 + ATP -> OAC | 48 | 3 | 41 | 56 |
| Mal -> Pyr + CO2 + NADPH | 5 | 3 | 0 | 11 |
| AKG + NADPH + NH3 -> Glu | 94 | 1 | 91 | 100 |
| Glu + ATP + NH3 -> Gln | 1 | 0 | NaN | 6 |
| Glu + ATP + 2*NADPH -> Pro | 0 | 0 | 0 | 3 |
| Glu + CO2 + Gln + Asp + AcCoA + 5*ATP + NADPH -> Arg + AKG + Fum + Ac | 0 | 0 | 0 | 4 |
| OAC + Glu -> Asp + AKG | 45 | 1 | 39 | 50 |
| Asp + 2*ATP + NH3 -> Asn | 0 | 0 | 0 | 7 |
| Pyr + Glu -> Ala + AKG | 0 | 0 | 0 | 15 |
| 3PG + Glu -> Ser + AKG + NADH | 0 | 0 | 0 | 4 |
| Ser <-> Gly + MEETHF | 0 | 0 | 0 | 3 |
| Gly <-> CO2 + MEETHF + NADH + NH3 | 0 | 0 | 0 | 1 |
| Thr <-> Gly + AcCoA + NADH | 0 | 0 | -3 | 0 |
| Ser + AcCoA + 3*ATP + 4*NADPH + SO4 -> Cys + Ac | 0 | 0 | 0 | 2 |
| Asp + Pyr + Glu + SucCoA + ATP + 2*NADPH -> LLDAP + AKG + Suc | 0 | 0 | 0 | 12 |

| | | | | |
|---|---|---|---|---|
| LLDAP -> Lys + CO2 | 0 | 0 | 0 | 12 |
| Asp + 2*ATP + 2*NADPH -> Thr | 45 | 1 | 34 | 47 |
| Asp + METHF + Cys + SucCoA + ATP + 2*NADPH -> Met + Pyr + Suc_int + NH3 | 0 | 0 | 0 | 2 |
| Pyr + Pyr + Glu + NADPH -> Val + CO2 + AKG | 0 | 0 | 0 | 8 |
| AcCoA + Pyr + Pyr + Glu + NADPH -> Leu + CO2 + CO2 + AKG + NADH | 0 | 0 | 0 | 2 |
| Thr + Pyr + Glu + NADPH -> Ile + CO2 + AKG + NH3 | 45 | 1 | 34 | 47 |
| PEP + PEP + E4P + Glu + ATP + NADPH -> Phe + CO2 + AKG | 3 | 1 | 0 | 4 |
| PEP + PEP + E4P + Glu + ATP + NADPH -> Tyr + CO2 + AKG + NADH | 0 | 0 | 0 | 4 |
| Ser + R5P + PEP + E4P + PEP + Gln + 3*ATP + NADPH -> Trp + CO2 + GAP + Pyr + Glu | 0 | 0 | 0 | 2 |
| R5P + FTHF + Gln + Asp + 5*ATP -> His + AKG + Fum + 2*NADH | 0 | 0 | 0 | 2 |
| MEETHF + NADH -> METHF | 0 | 0 | 0 | 2 |
| MEETHF -> FTHF + NADPH | 0 | 0 | 0 | 2 |
| NADH <-> NADPH | 75 | 5 | 65 | 86 |
| NH3.ext -> NH3 | 50 | 0 | 49 | 66 |
| SO4.ext -> SO4 | 0 | 0 | 0 | 2 |
| O2.ext -> O2 | 101 | 2 | 88 | 107 |
| CO2_unlabeled <-> CO2 | -93 | 63 | -95 | 21 |

| | | | | |
|---|---|---|---|---|
| CO2 -> CO2.ex | 0 | 63 | 0 | 49 |
| ATP -> ATP.maintenance | 1 | 0 | 1 | 1 |
| NADPH -> NADPH.maintenance | 1 | 0 | 1 | 1 |
| NADH + O2 -> 3*ATP | 27 | 3 | 22 | 52 |
| FADH2 + O2 -> 2*ATP | 61 | 4 | 27 | 68 |
| UQH2 + O2 -> 3*ATP | 13 | 0 | 13 | 13 |
| Suc -> Suc.ex | 2 | 0 | 2 | 4 |
| 2*Gln + 2*ATP + 2*FADH2 -> Indigo | 1 | 0 | 1 | 2 |
| 1.42*G6P + 0.56*F6P + 5.47*R5P + 2.77*E4P + 0.88*GAP + 8.86*3PG + 5.84*PEP + 14.35*Pyr + 20.49*AcCoA + 7.83*AKG + 11.04*OAC -> Biomass | 0.45 | 0 | 0.45 | 0.45 |
| SSR | 172 | | 128 | 198 |

**Table S5. Differentially Expressed Genes (DEGs) based on previous RNA-sequencing data collected in Banerjee et al., 2020**

| Locus | Unique DEGs for PSP (pTE327) compared to Eng (pTE219)* | Gene Name | Description |
|---|---|---|---|
| PP_2213 | 4.05 | - | acyl-CoA ligase |
| PP_2214 | 4.16 | *fadBA* | 3-hydroxyacyl-CoA dehydrogenase FadB2x |
| PP_2215 | 3.13 | - | 3-ketoacyl-CoA thiolase |
| PP_2216 | 2.81 | *acd* | acyl-CoA dehydrogenase family protein |
| PP_4116 | -2.57 | *aceA* | Isocitrate lyase |
| PP_4185 | -2.46 | *sucD* | succinyl-CoA synthetase, alpha subunit |
| PP_4186 | -2.64 | *sucC* | succinyl-CoA synthetase, beta subunit |

*DEGs were calculated as the Log2 ratios of 24 h RNA-sequencing data divided by 0 hr RNA-sequencing data for each strain, PSP and Eng, respectively. A fold change cutoff of 4 (|log2| >2) and a $p$-value < 0.001 was set as a cutoff to identify both large and statistically significant changes.

### Supplementary Methods

**Cellular Dry Weight Correlation.**

The cell dry weight (CDW) was measured over the course of growth and production in the WT and Engineered strains (**Fig. S1**). The WT CDW/$OD_{600}$ was constant throughout 24 hours and was correlated to 0.37 ± 0.02 mg/$OD_{600}$, consistent with previous measurements (Banerjee et al., 2020). Indigoidine production did not alter the Engineered strain CDW/$OD_{600}$ correlation during the early growth phase. After 6.5 hours, there was a decrease in the correlation, likely due to indigoidine absorption occurring around a similar wavelength as the OD wavelength measurement (indigoidine max absorbance occurred at a wave-length ($OD_{612}$, (Banerjee et al., 2020)). Overall, a CDW correlation of 0.37 ± 0.02 mg/$OD_{600}$ was used for WT strains and early growth phase indigoidine strains. A correlation of 0.29 ± 0.01 mg/$OD_{600}$ was used for indigoidine strains during the late production phase. After determining the cellular dry weight correlation, the strain performance was characterized under the cultivation conditions.

**Strain Scale-down.**

It was desirable to select culturing volumes that reduced experimental costs by minimizing labeled substrate use while still providing sufficient biomass for the analysis of low-abundance metabolites. Prior work had demonstrated that strain production was robust over a range of culture volumes (microplates, shaking flasks, and bioreactors) (Banerjee et al., 2020). The free headspace of the previous shaking flask experiments (60 mL liquid/250 mL flasks = 24%) was used as a guide to scale down the volume of liquid used in this study (14 mL liquid/50 mL shaking flasks = 28%).

**Flux Fitting.**

Amino acid fragment labeling data effects on the accuracy of the flux fits were assessed by analysis of the sum of square residues and the contribution of each fragment to the error as calculated in INCA  (Young, 2014). Redundant fragments and those whose inclusion lead to SSR fits falling outside the 95% confidence interval were removed from the fitting set. Additionally, fragments that led to standard errors larger than the best fit value were also removed. These fragments made up a minor portion of the data set and can be attributed to high signal-noise ratios. To further examine the flux value, Wuflux (He et al., 2016) was used to fit the 1,2-$^{13}$C glucose data as a cross validation, and generated similar flux maps.
